# Selective dysregulation of serotonin dynamics in the anterior cingulate cortex and central amygdala following binge alcohol consumption

**DOI:** 10.64898/2025.12.19.695484

**Authors:** Jobe L Ritchie, Maya R Eberle, Alison Roland, Lisa Taxier, Madison Campeau, Brianna George, Lili Kooyman, Thomas L Kash

## Abstract

Serotonin (5HT) is a critical modulator of brain function and behavior that is dysregulated in alcohol use disorder (AUD). The anterior cingulate cortex (ACC) and central nucleus of the amygdala (CeA) play distinct roles in AUD and undergo functional changes in 5HT signaling following binge drinking, but our understanding of real-time 5HT dynamics in these structures is lacking. We hypothesized that binge drinking would elicit brain-region specific dysfunction in 5HT dynamics during appetitive and aversive stimuli processing. Using fiber photometry with the GRAB5HT sensor, we identified distinct reward and aversion 5HT signaling motifs in the ACC and CeA. Consumption of alcohol and other tastants elicited a suppression of GRAB5HT signal in the ACC and an increase in 5HT signal in the CeA. In contrast, aversive stimuli similarly increased 5HT in both structures. The effect of binge drinking on 5HT function was surprisingly non-uniform, producing brain-region, sex-, and stimulus-specific dysfunction in 5HT signaling that dramatically shifted across weeks of alcohol experience. This suggests that adaptations in 5HT signaling are specific to neural circuits underlying discrete functions. Optogenetically stimulating 5HT terminals in the ACC and CeA increased avoidance behavior without being overtly rewarding or aversive, and stimulating 5HT release in the ACC blunted alcohol drinking. Together, these data identify distinct 5HT reward and aversion signaling motifs in the ACC and CeA and highlight early binge drinking as a critical stage of 5HT adaptation and a potential window for therapeutic intervention.

## Introduction

Alcohol misuse is associated with negative affective states, dysregulated reward processing, and a heightened risk for the development of an alcohol use disorder (AUD)^1,2^. Binge alcohol intoxication and subsequent withdrawal engage stress and reward circuitry, leading to neuronal dysfunction^3,4^, that contributes to maladaptive reward and aversion processing central to the pathophysiology of AUD^5^. Thus, developing a better understanding of the mechanisms underlying binge drinking-induced dysfunction is critical from a treatment perspective.

Serotonin (5-hydroxytryptamine, 5HT) is a neuromodulator central to reward and aversion processing, and affective regulation of behavior that is impacted in individuals with an AUD. Most of the 5HT in the brain is produced in the median and dorsal raphe nuclei (MRN and DRN, respectively). Clinical studies indicate that a history of AUD is associated with increased tryptophan hydroxylase, the rate limiting enzyme for 5HT production^6^, accompanied by altered 5HT transporter (SERT) and receptor binding in downstream targets^7–9^. These findings suggest that alcohol misuse induces lasting changes in 5HT circuitry.

Two brain regions impacted by AUD are the anterior cingulate cortex (ACC) and central nucleus of the amygdala (CeA). Problem drinkers exhibit blunted ACC recruitment during risky decision making^10^, and ACC functional connectivity predicts alcohol relapse risk^11^. Preclinically, binge alcohol consumption dynamically shifts ACC neuronal excitability based on the length of exposure^12^. Given this time-dependent effect, early versus prolonged binge drinking may elicit distinct adaptations in ACC, possibly driven by a shift in 5HT neuromodulation. Notably, these cortical alterations contrast with those in the amygdala, as SERT binding is increased in the ACC but decreased in the dorsal amygdala, including the CeA, in individuals with a history of AUD^7,9^. The CeA is a key anatomical substrate of both heightened negative affective states and excessive alcohol consumption^13^. Acutely, alcohol elicits 5HT release in the CeA to increase inhibitory neurotransmission^14^, which is impaired by chronic alcohol in a 5HT receptor dependent manner^15^. Together, these data suggest that alcohol misuse drives region-specific adaptations in 5HT signaling, which may underlie alcohol-induced deficits in reward and aversion processing. However, how binge drinking alters real-time 5HT dynamics in the ACC and CeA remains unknown.

This work aimed to elucidate the impact of binge drinking on 5HT dynamics in the ACC and CeA using *in vivo* fiber photometry and the genetically encoded serotonin sensor GRAB5HT. We measured 5HT signal during consummatory, avoidance, and aversive behaviors in alcohol-naïve mice or after the Drinking in the Dark (DID) model of binge drinking. We then combined genetically targeted anterograde tracing and optogenetics to assess the function of the MRN^sert^-ACC and DRN^sert^-CeA circuits in avoidance behavior and alcohol consumption. Together, these experiments reveal brain region-, sex-, and stimulus modality-dependent effects of binge drinking on 5HT dynamics and identify circuit-specific roles for ACC- and CeA-directed 5HT inputs in avoidance behavior and alcohol intake.

## Methods and Materials

### Subjects

Adult male and female mice (age 8-12 week) were singly housed on a 12:12 h reverse light-dark cycle (lights off at 7 AM) with *ad libitum* access to food (5P76 Irradiated RMH 3000 Rodent Diet) and water. For circuit tracing and optogenetic experiments, heterozygous *Sert^Cre^* mice (Strain# 014554, Jackson Laboratory) were bred in house (N = 20 males, 18 females; Fig. S1)^16^. For fiber photometry experiments, C57BL/6J mice (N = 27 males, 19 females) were sourced from Jackson Laboratories. All procedures were approved by the Institutional Animal Care and Use Committee of the University of North Carolina at Chapel Hill and performed in accordance with the National Institute of Health’s Guide for the Care and Use of Laboratory Animals.

### Surgery

Mice were anesthetized with 4% isoflurane and placed in a stereotaxic frame (Kopf Instruments). Craniotomies were performed above the injection site. For tracing and optogenetic experiments, AAV5-EF1A-DIO-eYFP-WPRE (EYFP control; Addgene; 3.6 x 10^12^) or AAV5-EF1A-DIO-hCHR2(H134R)-eYFP-WPRE (CHR2; Addgene; 3.6 x 10^12^) were infused at a rate of 100 nl/minute into the DRN (500 nl; AP: -4.5, ML: ±0.00, DV: −3.40, 20¼angle) or MRN (700 nl; AP: −4.25, ML: ±0.00, DV: −4.75, 10¼angle). The needle was left in place for 5 minutes to allow for diffusion. Fiber optic implants were placed bilaterally in the CeA (AP: −1, ML: ±2.5, DV: −4.70) or ACC (AP: 1.00, ML: ±0.30, DV: −1.60, 10¼angle), and secured in place using Metabond (Edgewood, New York). For fiber photometry experiments, 200 nl of a 1:5 cocktail of AAV8-hSyn-mCherry (Addgene; 3 x 10^12) and AAV8-hSyn-GRAB5HT3.0 (BrainVTA; 1.36 x 10^13) were injected into contralateral ACC and CeA using the above coordinates followed by fiber optic implantation, with mice counterbalanced for side placement. Experiments began 4-6 weeks post-surgery to allow for virus expression.

### Tissue Processing

Mice were anesthetized with 2.5% Tribromoethanol (1 ml, i.p.) and transcardially perfused with 0.01 M phosphate-buffered saline (PBS) followed by 4% paraformaldehyde (PFA) in PBS. Brains were extracted and post-fixed in 4% PFA overnight and then stored in PBS at 4 °C. 40 μm coronal sections were collected using a Leica VT1000S vibratome (Leica Microsystems) and stored in 0.02% Sodium Azide (Sigma Aldrich) in PBS.

### Fiber Photometry

Fiber photometry recordings were made using previously described methods^17,18^. Briefly, using a commercially available system from Neurophotometrics, Inc., GRAB5HT signal was recorded from the ACC and CeA simultaneously using a multi-branch patch cord (Doric, Québec, Canada). Pulses of 470 nm light (30 Hz, 50 μW) were bandpass filtered and focused onto a multi-branch patch cord by a 20x objective. Photometry data was processed using custom Matlab code. Traces were z-scored within session. Bout-associated traces were time-locked to start of behavior and normalized to pre-bout baseline. Video recordings were time locked to behavioral events using the open-source software Bonsai.

### Open Field Test

The open field test (OFT) was conducted in an empty plexiglass square arena with dimensions 50 cm x 50 cm x 40 cm illuminated by low intensity white LED light (100 lux). Mice were attached to fiber optic cables and allowed to freely explore the arena. The test continued for 10 (sensor experiment) or 12 (optogenetic experiments) minutes. For optogenetic experiments, mice received 20 Hz 470 nm laser stimulation (10 mW; SLOC, Shanghai China), in 3-minute blocks (Off/On x 2) signaled by Ethovision XT 14.0 (Noldus). Behavior was recorded with an overhead video camera and position was tracked using Bonsai or Ethovision software.

### Elevated Plus Maze

The elevated plus maze (EPM) assay was conducted on an elevated platform measuring 76 cm x 76 cm x 36 cm and consisting of two walled ‘closed’ arms and two unwalled ‘open’ arms. The maze was dimly illuminated from above by LED strips in a plus shape (open and closed arm lux, 25 and 15, respectively). Mice were attached to fiber optic cables and placed near the middle of the maze in the open arm with head oriented toward the center and allowed to freely explore. The test continued for 10 (sensor experiment) or 12 (optogenetic experiments) minutes. For optogenetic experiments, mice received 20 Hz 470 nm laser stimulation in 3-minute blocks (Off/On x 2). Behavior was recorded with an overhead video camera and position was tracked using Bonsai or Ethovision software. Accessory behaviors (head dip, guarded dip, rear, groom) were tracked using DeepLabCut (DLC)^19^ and Simple Behavioral Analysis (SimBA)^20^, as previously described^21^.

### Real-Time Place Preference

The real-time place preference assay (RTPP) was conducted over two days in an unmarked two-compartment rectangular apparatus measuring 52 cm x 26 cm x 26 cm. On day one, mice were allowed to freely explore the apparatus for 20 minutes. The following day, mice were allowed to explore the apparatus, and entry into one of the two compartments (counterbalanced) triggered 20 Hz 470 nm laser stimulation. Time spent on stimulation side over total time was used to calculate a preference score and compare between baseline and stimulation day.

### Drinking in the Dark (DID)

Mice were given free access to both water and 20% (v/v) alcohol bottles in the home cage three hours into the dark cycle for 2 h on Monday through Wednesday and 4 h on Thursdays. Alcohol and water bottle positions were swapped on alternating days to account for side preference and drip values from bottles on empty cages were subtracted when calculating daily consumption. This schedule continued for 4 weeks. Blood ethanol concentrations were determined from tail blood collected following the final 4 h DID session using the AM1 Analox Analyzer (Analox Instruments) and used to confirm binge levels of intoxication of approximately 80 g/dl.

### Consummatory Behaviors with Fiber Photometry

Testing was conducted under dim red light in a plastic cage with a rimmed top beginning at approximately 10 AM. Mice were water restricted for 3-6 hours prior to testing. Following 5 minutes of acclimation, the mice were allowed to consume 2-3 bouts each of 20% alcohol (v/v), 3% sucrose in water, and high fat diet (HFD, Research Diets, 60 kcal% fat), in that order. The minimum bout duration was set at 1 second. Testing was concluded when all bouts were achieved or if the mice did not complete all bouts within one hour (n = 5 sessions).

### Anterograde 5HT Circuit Tracing

Brain tissue from male Sert-cre mice with DIO-EYFP expression restricted to either DRN or MRN (N = 3 per region) was used for anterograde tracing. EYFP fluorescence was enhanced using rabbit anti-GFP conjugated to 488 incubated overnight at room temperature (1:1000; Jackson Immunoresearch). Regions of interest were imaged (2-4 images/region/subject) on an Echo confocal microscope (San Diego, CA) at 10x magnification. Images were background subtracted, and mean pixel intensity was quantified in FIJI software^22^. Values were normalized within brain region across samples.

### Home Cage Alcohol Drinking with Optogenetics

Optogenetic stimulation during alcohol consumption testing was conducted in the home cage in the dark. Mice were acclimated to drinking 20% alcohol solution while tethered for four 1-hour sessions prior to testing. Mice were then allowed free access to 20% alcohol solution for 1 hour during a baseline session (no laser stimulation) and stimulation session, with counterbalanced order across two days. During the stimulation session, mice received 20 Hz 470 nm laser stimulation in 5-minute blocks (Off/On x 6). Bottles were weighed before and after the sessions to calculate EtOH consumption. Empty cage drip values were subtracted from total volume.

### Home Cage Behaviors, Sucrose Splash, and Scruff Assays

Mice were attached to fiber optic cables and allowed to freely explore their home cage with the lid removed under bright illumination (200 lux). Behavior was recorded using an overhead camera. The first 5 minutes of the session were used to assess home cage behaviors. A 10% sucrose solution was then applied to the back of the mouse using a spray bottle. The following 10 minutes were analyzed for post-splash behaviors. The mice were then scruffed for 1 minute, followed by 3 minutes of post-scruff observation. Behaviors (rear, groom, dig) were tracked using DeepLabCut (DLC)^19^ and Simple Behavioral Analysis (SimBA)^20^, as previously described^21^.

### Slice Electrophysiology

Whole-cell patch clamp recordings were performed as previously described^23^ in acute coronal brain slices (250 µM) containing the DRN and MRN of Sert^cre^ mice expressing AAV5-EF1A-DIO-hCHR2(H134R)-eYFP-WPRE. Slices were prepared in ice-cold cutting solution and recovered in oxygenated aCSF at 35 °C. eYFP-labeled neurons were recorded in current clamp using a potassium gluconate internal solution. Signals were acquired with a Multiclamp 700B amplifier, filtered at 3 kHz, digitized at 10 kHz, and analyzed offline. Action potentials were evoked by 1 Hz, 1 ms pulses of blue light (490 nm).

### Statistical Analysis

Data from misplaced virus expression or fiber implants were excluded from analysis on a per region basis (n = 3 ACC and 10 CeA exclusions from 13 mice). Statistical analyses were performed using GraphPad Prism or SPSS software. Data were analyzed using Analysis of Variance (ANOVA) or mixed factorial ANOVA, with sex (male, female), group (Water, DID), virus (EYFP, CHR2), as between-subjects factors and time/day as within-subjects factors, where appropriate. Planned comparison t-tests were used where appropriate. Significant effects were followed up using Tukey’s *post hoc* test. Significance was set at *p* = 0.05.

## Results

### Males exhibit more robust changes in 5HT in a novel anxiogenic environment than females

Prior literature employing raphe lesion or genetic knockdown of *Sert* suggests that 5HT signaling regulates exploration of a novel environment, with distinct roles for MRN and DRN^24,25^. However, potential sex- and region-specific differences in 5HT dynamics have not been assessed. To this end, *in vivo* photometry recordings of mice expressing the 5HT sensor GRAB5HT in the ACC and CeA were acquired during free exploration using the open field test (OFT; **Fig. 1A)** prior to alcohol drinking. Behaviorally, females spent less time in the center of the OFT and more time in the corner, but had a similar number of center entries compared to males **(Fig. 1D-E)**. This behavioral difference was accompanied by sex differences in 5HT activity during exploration. In the ACC, males exhibited 5HT signal that increased in the center relative to the corner, an effect not seen in females **(Fig. 1F-G)**. CeA 5HT signal in the center of the OFT increased relative to the corner for both sexes but was higher for males than females **(Fig. 1H-F)**. Together, these data indicate that in male mice, 5HT signaling has a greater dynamic range to exploration of the anxiogenic zone of a novel environment.

**Figure 1.**
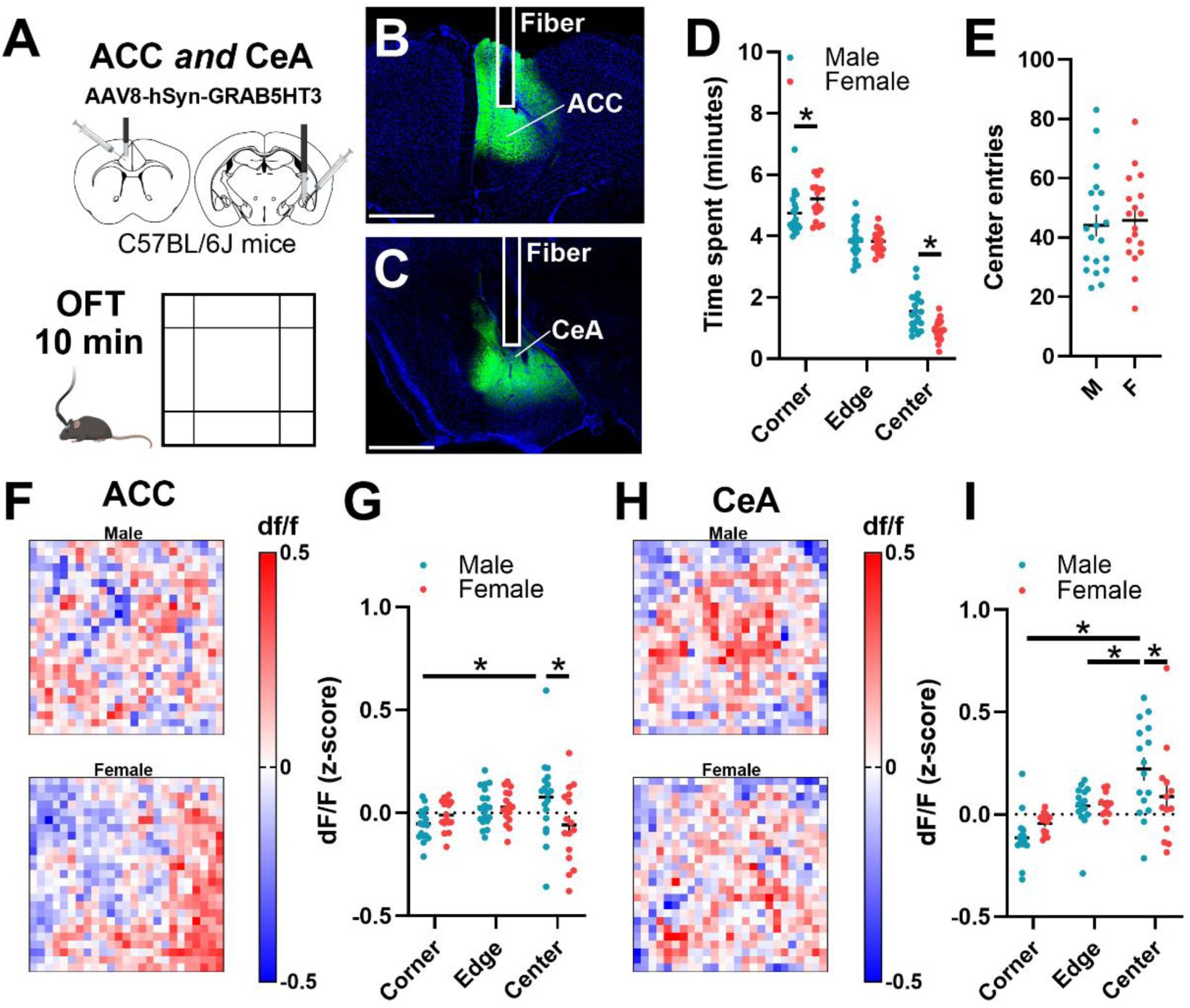
Increased male 5HT signal in anxiogenic zone of novel context. **(A)** Experimental design. Male (n = 20) and female (n = 18) C57BL/6J mice with GRAB5HT and fiber optic implants directed at the anterior cingulate cortex and central nucleus of the amygdala were allowed to freely explore a novel open field (OFT) for 10 minutes. **(B,C)** Representative examples of virus expression and fiber placement in **(B)** anterior cingulate cortex and **(C)** central nucleus of the amygdala. **(D)** Females spent more time in the corner and less time in the anxiogenic center of the open field (2-way ANOVA, sex x zone interaction, F_(2,108)_ = 9.12, *p* = 0.0002; Sidak’s post hoc tests, female corner > male corner *p* = 0.03, female center < male center, *p* = 0.003). **(E)** Number of center entries for males and females. **(F)** Representative heatmaps of 5HT signal in the ACC for (upper) males and (lower) females. **(G)** Quantification of average ACC 5HT signal in each zone revealed that male signal increased in the center relative to the corner and that female signal in center was lower than males (2-way ANOVA, sex x zone interaction, F_(2,99)_ = 4.69, *p* = 0.01; Sidak’s post hoc tests, male center > male corner, *p* = 0.006, female center < male center *p* = 0.002). **(H)** Representative heatmaps of 5HT signal in the CeA for (upper) males and (lower) females. **(I)** Quantification of average CeA 5HT signal in each zone revealed that male signal increased in the center and edge relative to the corner and that female signal in center was lower than males (2-way ANOVA, sex x zone interaction, F_(2,81)_ = 3.49, *p* = 0.04; Sidak’s post hoc tests, male center > male corner, *p* < 0.001, male center > male edge, *p* = 0.003, female center < male center *p* = 0.02). Abbreviations: anterior cingulate cortex (ACC); basolateral amygdala (BLA); central nucleus of the amygdala (CeA).

### A history of binge drinking blunts 5HT response in the ACC to alcohol consumption and has sex-specific effects on sucrose and HFD associated 5HT signal

The effects of binge drinking on neuronal excitability vary depending on brain region and period of intake^3^. However, shifts in real-time 5HT dynamics across different periods of binge drinking is not known. After the OFT, we assessed the 5HT response in the ACC and CeA of alcohol-naïve mice to consumption of 20% alcohol, 3% sucrose, and high fat diet (HFD; **Fig. 2A-C)**. Mice then underwent 4 weeks of alcohol drinking using the Drinking in the Dark (DID) model, with additional photometry recording sessions after the first and fourth weeks. During DID, females drank more alcohol than males but achieved similar blood alcohol concentrations during the final DID session **(Fig. S2B-C)**. In the ACC, 5HT release was typically reduced during consummatory behaviors. Following one week of binge drinking, the ACC 5HT response to alcohol consumption was blunted in both sexes **(Fig. 2E, Q)**, an effect that was no longer observed after 4 weeks **(Fig. 2F, R)**. Interestingly, the 5HT response to sucrose in the ACC was elevated at the 1-week time point in water control male mice only **(Fig. 2I)**, while DID mice showed no change across weeks for males or females. DID males also showed a trend towards decreased 5HT response to high fat diet (HFD) following DID that was absent in females **(Fig. 2M-O)**. Thus, males and females exhibited similar shifts in the ACC 5HT response to alcohol consumption, but males showed an augmented 5HT decrease in response to other tastants.

**Figure 2.**
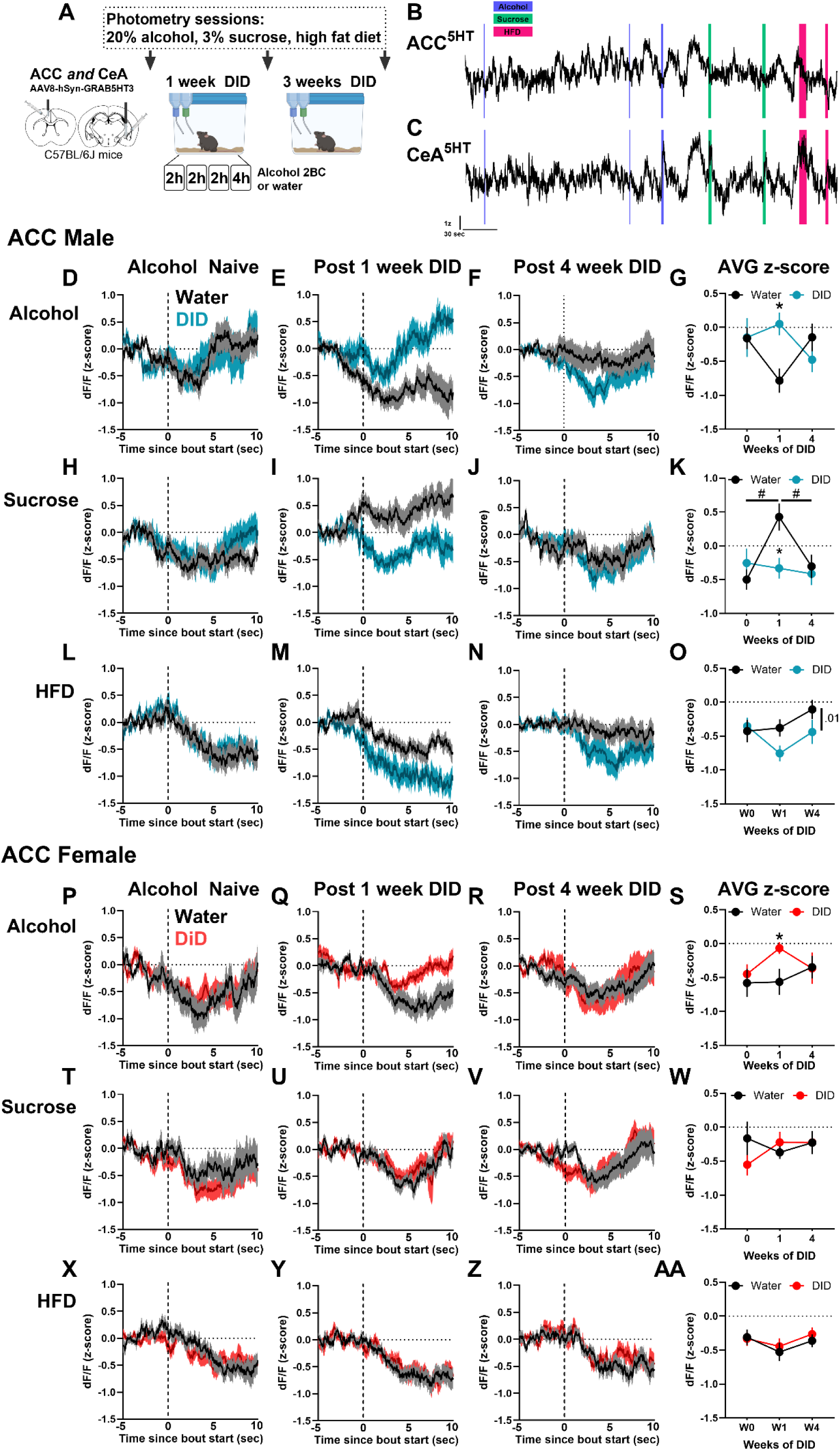
Sex-specific effects of binge drinking on 5HT response to consummatory behaviors in ACC. **(A)** Experimental design. Male and female C57BL/6J mice received GRAB5HT and fiber optic implants directed at the anterior cingulate cortex and central nucleus of the amygdala and underwent 4 weeks of 2-bottle choice Drinking in the Dark (DID) binge drinking of 20% alcohol (N = 10 male, 10 female) or served as water controls (N = 10 male, 9 female). Photometry recording sessions to assess *in vivo* 5HT dynamics while consuming 20% alcohol, 3% sucrose, and high fat diet (HFD) were conducted in alcohol-naïve mice and after 1 and 4 weeks of DID. **(B)** Representative GRAB5HT photometry traces from the ACC and **(C)** CeA of an alcohol naïve mouse during home cage consummatory behavior recording. Blue, green, and pink highlighted sections represent bouts of alcohol, sucrose, and HFD consumption, respectively. **Male ACC:** 5HT response in ACC during consumption of alcohol in water control and DID mice after **(D)** 0 weeks, **(E)** 1 week, and **(F)** 4 weeks of DID. The dotted line represents the start of a bout. **(G)** Mean alcohol bout z-score across weeks. DID mice exhibited a significantly higher ACC 5HT response to alcohol consumption than water controls after 1 week of DID (2-way ANOVA, day x group interaction, F_(2,83)_ = 4.44, p = 0.01, Sidak’s post hoc, 1 week DID > 1 week water, p = 0.005). 5HT response in ACC during consumption of sucrose after **(H)** 0 weeks, **(I)** 1 week, and **(J)** 4 weeks of DID. **(K)** Mean sucrose bout z-score across weeks. DID mice exhibited a significantly lower ACC 5HT response to sucrose consumption than water controls after 1 week of DID (2-way ANOVA, day x group interaction, F_(2,89)_ = 4.26, *p* = 0.02, Sidak’s post hoc, 1 week DID > 1 week water, *p* = 0.003). Additionally, male water mice 5HT response to sucrose was higher at the 1 week time point compared to other time points (week 1 > weeks 0 and 4, p < 0.05). 5HT response in ACC during consumption of HFD after **(L)** 0 weeks, **(M)** 1 week, and **(N)** 4 weeks of DID. **(O)** Mean HFD bout z-score across weeks. DID mice exhibited a trend towards lower ACC 5HT response to HFD consumption compared to water controls (2-way ANOVA, group main effect, F_(1,90)_ = 2.93, *p* = 0.09). **Female ACC:** 5HT response in ACC during consumption of alcohol in water control and DID mice after **(P)** 0 weeks, **(Q)** 1 week, and **(R)** 4 weeks of DID. **(S)** Mean alcohol bout z-score across weeks. DID mice exhibited a significantly higher ACC 5HT response to alcohol consumption than water controls after 1 week of DID (Planned comparison Sidak’s post hoc,1 week DID > 1 week water, *p* = 0.05). 5HT response in ACC during consumption of sucrose in water control and DID mice after **(T)** 0 weeks, **(U)** 1 week, and **(V)** 4 weeks of DID. **(W)** Mean sucrose bout z-score across weeks. 5HT response in ACC during consumption of HFD in water control and DID mice after **(X)** 0 weeks, **(Y)** 1 week, and **(Z)** 4 weeks of DID. **(AA)** Mean HFD bout z-score across weeks.

### Binge drinking augments 5HT response in the CeA to alcohol consumption and preserves sucrose salience in males

In contrast to the ACC, the CeA exhibited an increase in GRAB5HT signal during consummatory behaviors, with blunted or absent responses in females **(Fig. 3)**. DID augmented the CeA 5HT response to alcohol consumption in males **(Fig. 3A-D)**, but not females **(Fig. 3M-P)**. Male DID mice showed a stable 5HT response to sucrose consumption across weeks that was surprisingly absent in water mice at the 4-week time point **(Fig. 3E-G)**. In contrast, female mice did not exhibit significant changes in 5HT dynamics across four weeks of binge drinking **(Fig. 3Q-T)**. The 5HT response in the CeA to HFD remained stable across weeks, independent of sex **(Fig. 3I-L, 3U-X)**. Together, these data suggest that DID enhances or maintains CeA 5HT response to alcohol and sucrose in males and that females are resilient to dysregulation by DID.

**Figure 3.**
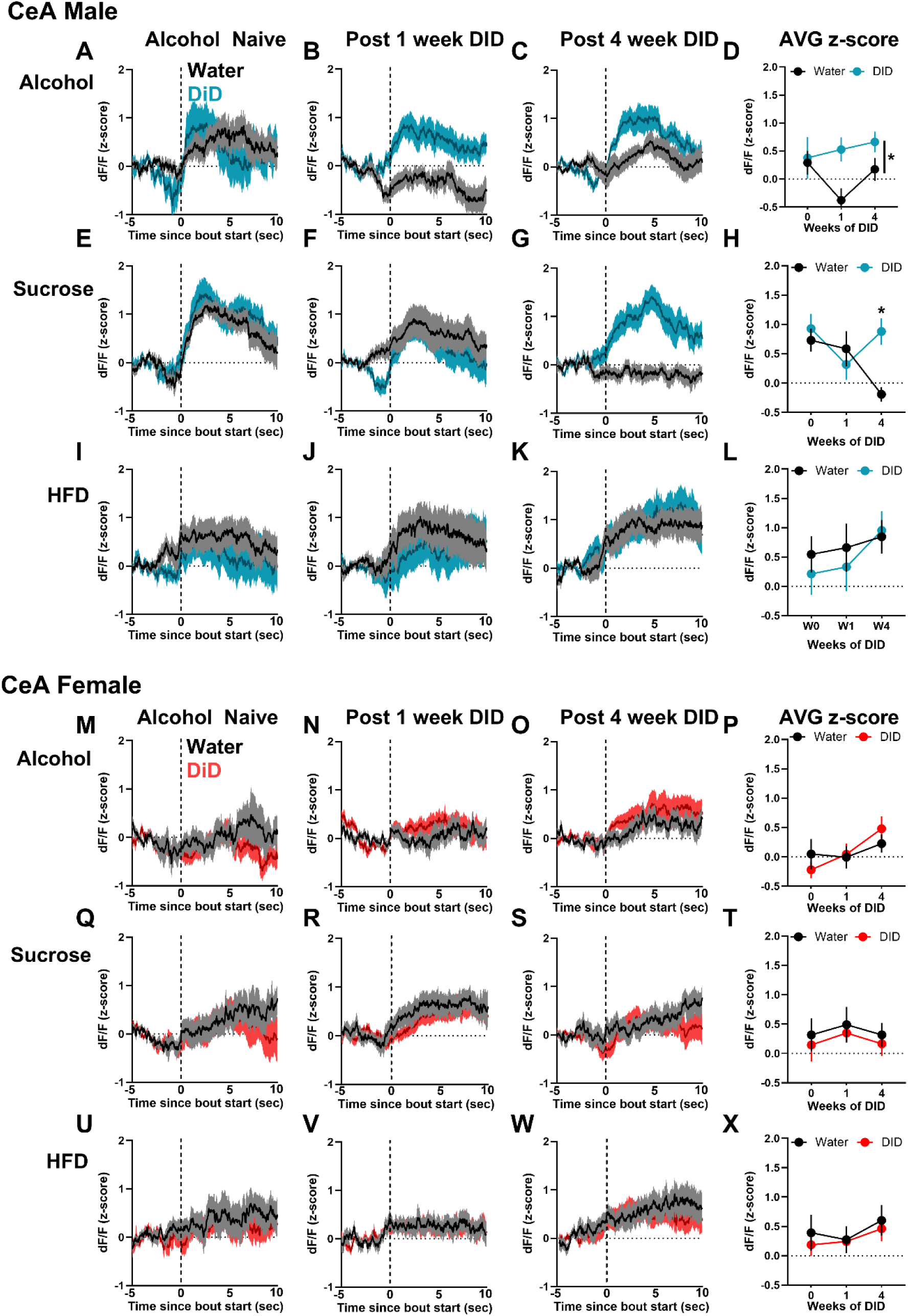
Binge drinking augments 5HT response in the CeA to alcohol and sucrose consumption in males. Male and female C57BL/6J mice received GRAB5HT and fiber optic implants directed at the anterior cingulate cortex and central nucleus of the amygdala and underwent 4 weeks of 2-bottle choice Drinking in the Dark (DID) binge drinking of 20% alcohol (N = 10 male, 10 female) or served as water controls (N = 10 male, 9 female). Photometry recording sessions to assess in vivo 5HT dynamics while consuming 20% alcohol, 3% sucrose, and high fat diet (HFD) were conducted in alcohol-naïve mice and after 1 and 4 weeks of DID. See prior figure for experimental timeline. **Male CeA:** 5HT response in CeA during consumption of alcohol in water control and DID mice after **(A)** 0 weeks, **(B)** 1 week, and **(C)** 4 weeks of DID. The dotted line represents the start of a bout. **(D)** Mean alcohol bout z-score across weeks. DID mice exhibited a significantly higher ACC 5HT response to alcohol consumption than water controls (2-way ANOVA, group main effect, F_(1,66)_ = 6.97, *p* = 0.01, planned comparison Sidak’s post hoc, 1 week DID > 1 week water, *p* = 0.004). 5HT response in CeA during consumption of sucrose in water control and DID mice after **(E)** 0 weeks, **(F)** 1 week, and **(G)** 4 weeks of DID. **(H)** Mean sucrose bout z-score across weeks. DID mice exhibited a significantly higher ACC 5HT response to sucrose consumption than water controls after 4 weeks of DID (2-way ANOVA, group x day interaction, F_(2,69)_ = 6.97, *p* = 0.03, Sidak’s post hoc, 4 week DID > 4 week water, *p* = 0.004). 5HT response to sucrose consumption decreased across time in water control mice (water week 1 > water week 4, *p* = 0.04). 5HT response in CeA during consumption of HFD in water control and DID mice after **(I)** 0 weeks, **(J)** 1 week, and **(K)** 4 weeks of DID. **(L)** Mean HFD bout z-score across weeks. **Female CeA:** 5HT response in CeA during consumption of alcohol in water control and DID mice after **(M)** 0 weeks, **(N)** 1 week, and **(O)** 4 weeks of DID. **(P)** Mean alcohol bout z-score across weeks. 5HT response in CeA during consumption of sucrose in water control and DID mice after **(Q)** 0 weeks, **(R)** 1 week, and **(S)** 4 weeks of DID. **(T)** Mean sucrose bout z-score across weeks. 5HT response in CeA during consumption of HFD in water control and DID mice after **(U)** 0 weeks, **(V)** 1 week, and **(W)** 4 weeks of DID. **(X)** Mean HFD bout z-score across weeks.

### 5HT signal in EPM scales to aversion, with mixed effects of DID on anxiety-like and stress coping behaviors and associated 5HT responses

Four to five days following 4 weeks of DID, mice were assessed for avoidance behaviors and 5HT dynamics in the elevated plus maze (EPM; **Fig. 4A)**. Male DID mice showed fewer open arm entries and distance traveled relative to water males, with no difference in females **(Fig. 4B-C)**. There was no effect of DID on time spent in zones of EPM **(Fig. S3A)**. 5HT signal in the ACC and CeA exhibited a clear distinction between open, center, and closed zones of the EPM that scaled with the aversiveness of the zones **(Fig. 4D-G)**. Averaging ACC 5HT signal by zone revealed a blunting of the response in DID females only **(Fig. 4H)**. Time spent in all zones negatively correlated with 5HT signal in both regions **(Fig. 4I, 4K)**. However, when analyzing time spent in the closed arm, a positive correlation is observed. Analysis of 5HT signal during initiation of exploration and retreat from the open arm revealed that male DID mice showed an elevated ACC 5HT response relative to water controls that was absent in females and did not similarly occur in the CeA **(Fig. 4L, 4N)**. 5HT signal during initiation of explore/retreat bouts did not correlate with locomotor velocity **(Fig. S3B-C)**. Notably, the likelihood to explore the open arm decreased with increased 5HT signaling, while the likelihood to retreat from the open arm increased, independent of brain region **(Fig. 4M, 4O)**. Additional EPM behaviors (head dip, guarded dip, rear, groom) were analyzed using machine learning; no DID effect was observed on behavior count, but DID blunted CeA 5HT signal during exploratory behaviors in a sex dependent manner **(Fig. S4A-D)**. Overall, these data suggest that elevated 5HT drives avoidance behavior and promotes retreat. In contrast to consummatory behaviors, where the ACC and CeA showed opposite 5HT responses, the direction of response was similar in both regions during avoidance behavior, and DID blunted 5HT signaling in a sex- and behavior-specific manner.

**Figure 4.**
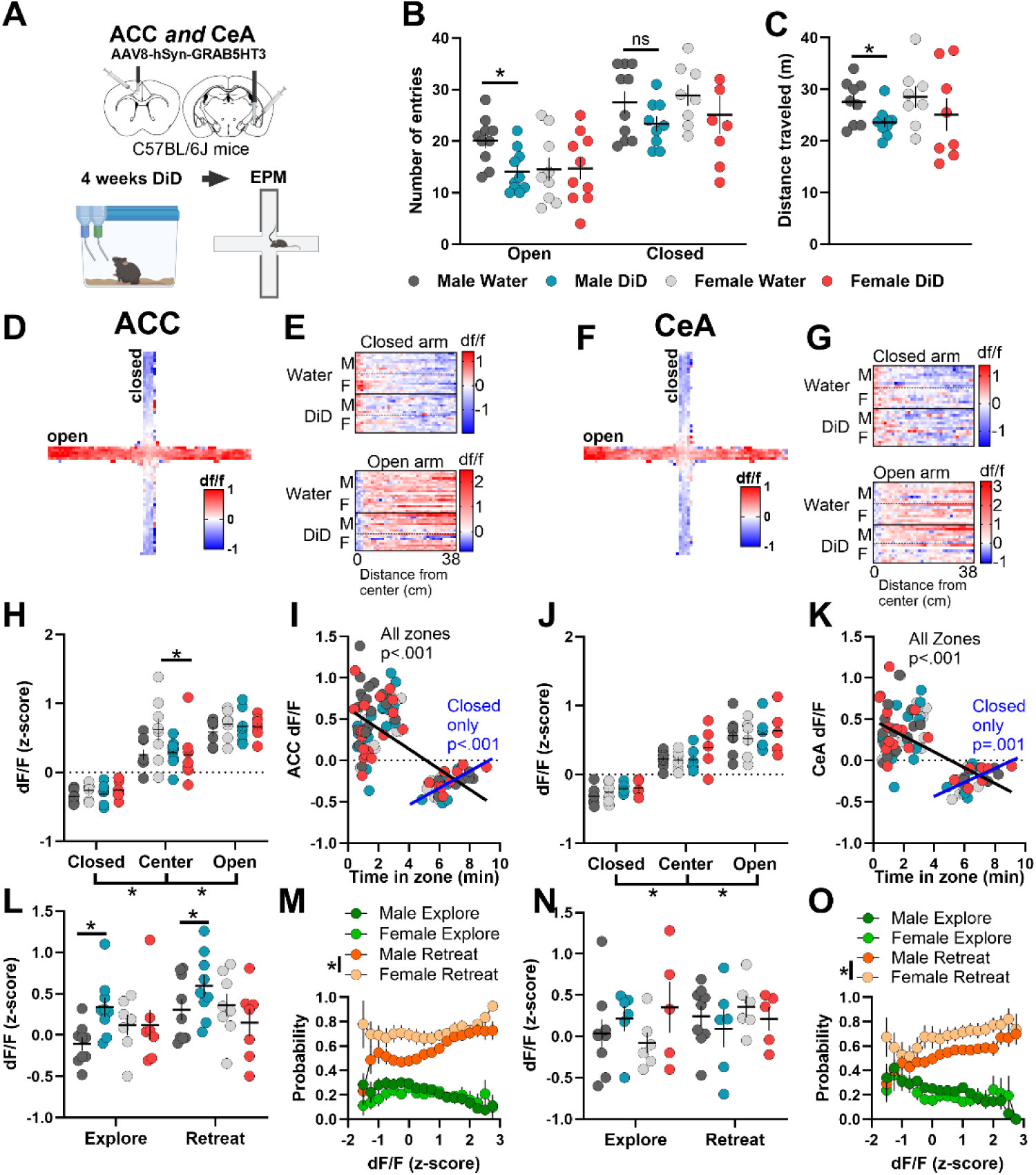
Binge drinking increases avoidance behaviors in males and 5HT signal in the ACC and CeA convey ambiguous threat. **(A)** Experimental timeline. Male and female C57BL/6J mice received GRAB5HT and fiber optic implants directed at the anterior cingulate cortex and central nucleus of the amygdala and underwent 4 weeks of 2-bottle choice Drinking in the Dark (DID) binge drinking of 20% alcohol (N = 10 male, 10 female) or served as water controls (N = 10 male, 9 female). Mice were allowed to explore an elevated plus maze (EPM) for 10 minutes 5 days after their final DID session. **(B)** Total open and closed arm entries. Male DID mice had fewer open arm entries than male water controls (Welch corrected t-test, t_(17.9)_ = 3.09, *p* = 0.006) but not closed arm entries (*p* = 0.13). **(C)** Distance traveled in the EPM. Male DID mice traveled less distance than male water mice (Welch corrected t-test, t_(13.9)_ = 2.33, *p* = 0.04). **(D)** Representative heatmap of ACC 5HT signal in the EPM. **(E)** Per subject heatmap of distance from center in (upper) closed and (lower) open arm. **(F)** Representative heatmap of CeA 5HT signal in the EPM. **(G)** Per subject heatmap of distance from center in (upper) closed and (lower) open arm. **(H)** Average z-scored ACC 5HT signal by zone of the EPM. DID females had blunted 5HT signal in center of the EPM (3-way ANOVA, sex x group interaction, F_(1,29)_ = 4.07, *p* = 0.05, Sidak’s post hoc, DID female center < water female center, *p* = 0.04). **(I)** Correlational analysis of ACC 5HT signal and total time in each zone. ACC 5HT signal negatively correlated with time when including all zones (Pearson correlation, r = -0.67, *p* < 0.0001) but positively correlated with time in closed arm (r = 0.62, *p* < 0.0001). **(J)** Average z-scored CeA 5HT signal by zone of the EPM. CeA 5HT signal varied by zone only with increased signal in center and open arm compared to closed arm (3-way ANOVA, zone main effect, F_(2,44)_ = 104.79, *p* < 0.001, Sidak’s post hoc, closed < center < open, *p* < 0.001). **(K)** Correlational analysis of CeA 5HT signal and total time in each zone. CeA 5HT signal negatively correlated with time when including all zones (Pearson correlation, r = -0.63, *p* < 0.0001) but positively correlated with time in closed arm (r = 0.55, *p* = 0.001). **(L)** Average ACC 5HT signal during initiation of exploration and retreat from open arm. Male 5HT signal was higher during retreat, and male DID mice 5HT signal was higher than male water mice (2-way ANOVA, group main effect, F_(1,32)_ = 9.92, *p* = 0.004, behavior main effect, F_(1,32)_ = 8.34, *p* = 0.007). **(M)** Probability to explore or retreat from open arm in subsequent 10 seconds based on ACC 5HT signal. **(N)** Average CeA 5HT signal during initiation of exploration and retreat from open arm. **(O)** Probability to explore or retreat from open arm in subsequent 10 seconds based on CeA 5HT signal.

### DID induced changes in 5HT response to aversive stimuli in males and females

Binge drinking has sex-specific effects on behavioral and cellular responses to aversive stimuli^26^, but the potential role of changes in 5HT dynamics is not known. Thus, in the same group of mice, we next assessed the effects of 4 weeks of binge drinking on spontaneous home cage behaviors and 5HT dynamics during non-noxious aversive stimuli **(Fig. 5A)**. Analysis of home cage behaviors did not reveal effects of DID in the present study **(Fig. S5A-I)**. The non-noxious aversive stimulus of sucrose splash produced a robust increase in 5HT signal in the ACC, while a 1-minute scruff increased 5HT signal in both the ACC and CeA. In the ACC, female DID mice showed an augmented 5HT response to sucrose splash but not scruff, with no effect of DID in males **(Fig. 5B-G)**. Contrastingly, in the CeA, male DID mice showed a blunted 5HT response to scruffing, with no effect of DID in females, and minimal response to sucrose splash in either sex **(Fig. 5H-M)**. These data highlight sex differences in the effect of DID on aversive processing and suggest that stimulus modality is a critical factor.

**Figure 5.**
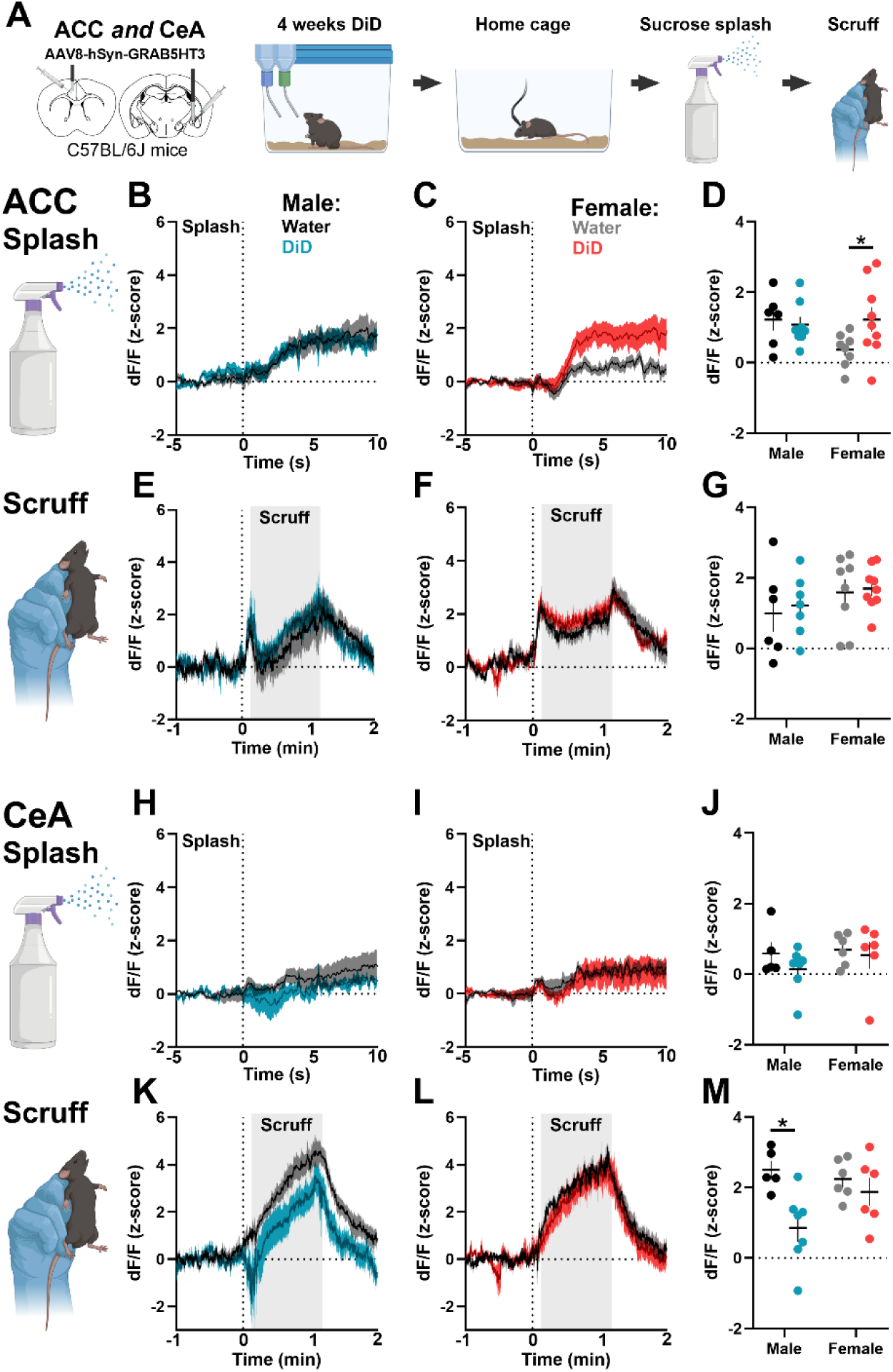
Binge drinking alters 5HT response to aversive stimuli in a sex- and region-dependent manner. **(A)** Experimental timeline. Male and female C57BL/6J mice received GRAB5HT and fiber optic implants directed at anterior cingulate cortex and central nucleus of the amygdala and underwent 4 weeks of 2-bottle choice Drinking in the Dark (DID) binge drinking of 20% alcohol (N = 10 male, 10 female) or served as water controls (N = 10 male, 9 female). Mice were allowed to explore their home cage for 5 minutes then received a spray of 10% sucrose solution directed at their back. Ten minutes later, mice were scruffed for 1 minute and then allowed to explore their home cage for 2 minutes. ACC 5HT response following sucrose splash in **(B)** males and **(C)** females. **(D)** Average ACC 5HT signal following splash. DID females had augmented 5HT response to sucrose splash compared to water females (Welch’s t-test, t_(11.22)_ = 2.18, *p* = 0.05). ACC 5HT response to scruff in **(E)** males and **(F)** females. **(G)** Average ACC 5HT signal following splash. CeA 5HT response following sucrose splash in **(H)** males and **(I)** females. **(J)** Average CeA 5HT signal following splash. CeA 5HT response to scruff in **(K)** males and **(L**) females. **(M)** Average CeA 5HT signal following scruff. DID males had augmented 5HT response to scruff compared to water males (Welch’s t-test, t_(9.64)_ = 3.52, *p* = 0.006).

### 5HT outputs of the MRN and DRN include the ACC and CeA, respectively

There is an inconsistent and limited literature on MRN and DRN 5HT output targets of 5HT to AUD-associated structures^27,28^. To inform brain region targeting in subsequent experiments, we used Sert-cre mice to genetically express EYFP in 5HT neurons **(Fig. S1A-H)** in the MRN or DRN and then quantified process labeling in downstream targets **(Fig. 6A-E)**. Process labeling of MRN^Sert^ and DRN^Sert^ revealed distinct projection patterns, shown organized by the primary input region **(Fig. 6F)**. Notably, the ACC and CeA fell into the MRN^Sert^- and DRN^Sert^-dominant categories, respectively, and these circuits were investigated in subsequent experiments.

**Figure 6.**
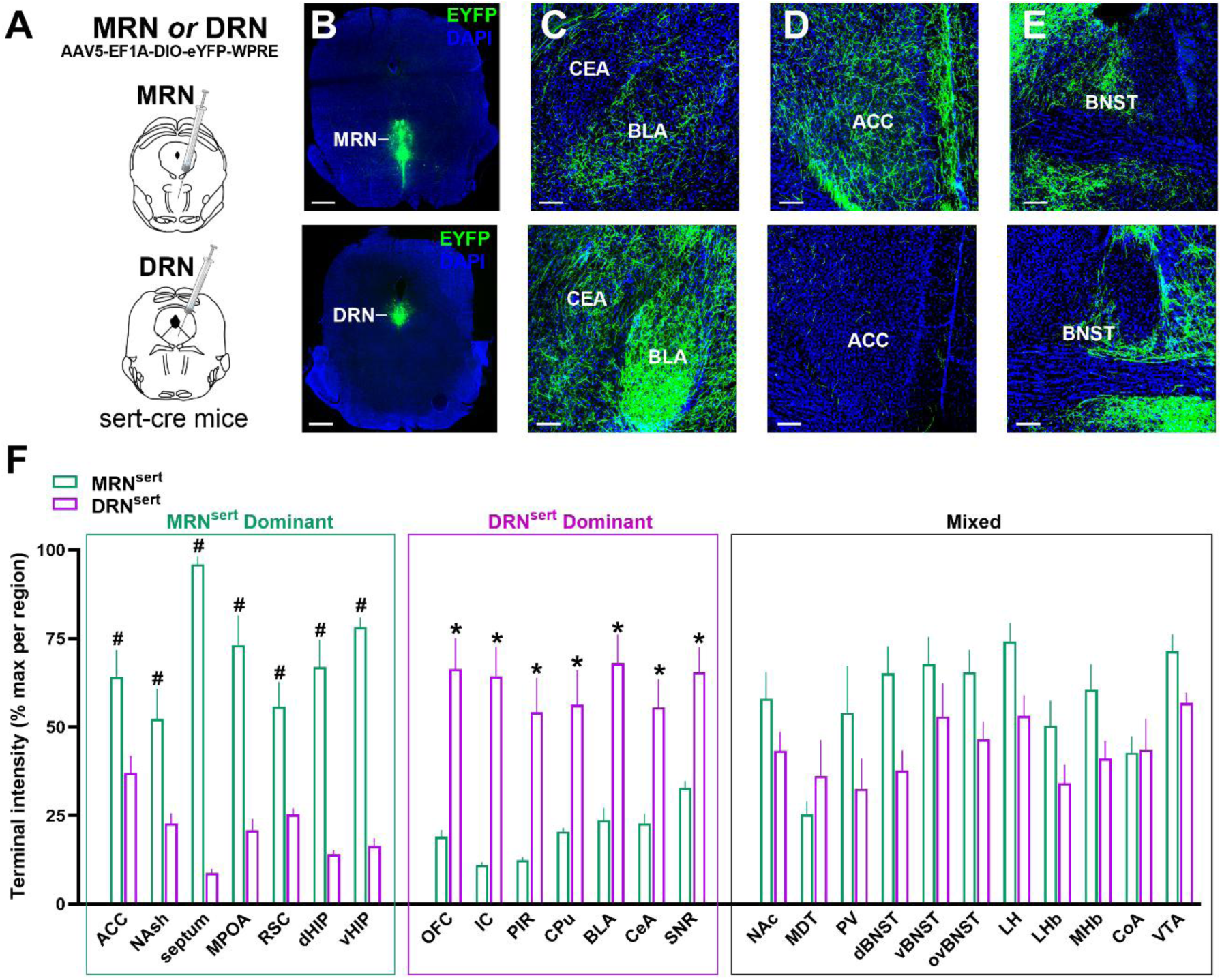
Distinct output targets of MRN^Sert^ and DR^Sert^. **(A)** Experimental design. Sert-cre mice received a cre-dependent EYFP virus in the MRN or DRN (n = 3 males per brain region) to label 5HT neurons in **(B, upper)** MRN and **(B, lower)** DRN. Process labeling was enhanced with a 488-congugated anti-GFP antibody and imaged in 2-4 replicates per subregion per subject. Representative example images of process labeling in **(C)** amygdala complex **(D)** anterior cingulate cortex and **(E)** bed nucleus of the stria terminalis originating from (upper panels) MRN and (lower panels) DRN. **(F)** Quantification of percent max terminal intensity revealed distinct output targets between groups (2-way ANOVA, group x region interaction, F_(24,445)_ = 17.00, *p* < 0.001). Brain regions were organized into MRN^Sert^-dominant (^#^) or DR^Sert^-dominant (*) based on significant post hoc tests (Sidak’s test, *p* < 0.05). Brain regions that DID not reach significance fell into the Mixed category. Abbreviations: anterior cingulate cortex (ACC); nucleus accumbens shell (NAsh); medial preoptic area (MPOA); retrosplenial cortex (RSC); dorsal hippocampus (dHIP); ventral hippocampus (vHIP); orbitofrontal cortex (OFC); insular cortex (IC); piriform cortex (PIR); caudate putamen (CPu); basolateral amygdala (BLA); central nucleus of the amygdala (CeA); substantia nigra reticulata (SNR); nucleus accumbens core (NAc); mediodorsal thalamus (MDT); paraventricular thalamus (PV); dorsolateral bed nucleus of the stria terminalis (dBNST); vental bed nuleus of the stria terminals (vBNST); oval nucleus of the bed nucleus of the stria terminals (ovBNST); lateral hypothalamus (LH); lateral habenula (LHb); medial habenula (MHb); cortical amygdala (CoA); ventral tegmental area (VTA).

### 5HT terminal stimulation in the ACC and CeA is anxiogenic and circuit-dependently blunts alcohol drinking

The ACC and CeA receive 5HT input primarily from MRN and DRN, respectively, but the role of MRN^Sert^-ACC and DRN^Sert^-CeA circuits in avoidance behavior and alcohol consumption have not been assessed. Thus, we used genetically targeted optogenetics in Sert-cre mice to stimulate terminals in the MRN^Sert^-ACC or DRN^Sert^-CeA circuit **(Fig. 7A-C)**. Stimulation of 5HT terminals in the ACC and CeA similarly resulted in less center time in the OFT, without altering distance traveled, indicating an anxiogenic effect **(Fig. 7D-H)**. However, neither manipulation produced a conditioned place preference or avoidance in the Real Time Place Preference (RTPP) assay, suggesting the signal is not robustly valenced or the effects of stimulation are context dependent **(Fig. 7I-M)**. After 4 days of acclimation to drinking 20% alcohol while tethered, we assessed home cage drinking on a baseline (no-stimulation) and stimulation day (5 minutes on/5 minutes off, 20hz). Stimulation of 5HT terminals in the ACC decreased alcohol consumption relative to baseline in CHR2, but not EYFP mice **(Fig. 7N-P)**, while stimulation directed at 5HT processes in the CeA had no effect on drinking **(Fig. 7Q-R)**. Thus, ACC and CeA 5HT signaling may play similar roles in avoidance behavior but diverge in their role in regulation of alcohol consumption, mirroring the distinct reward and aversion 5HT signaling motifs observed in our 5HT sensor experiments.

**Figure 7.**
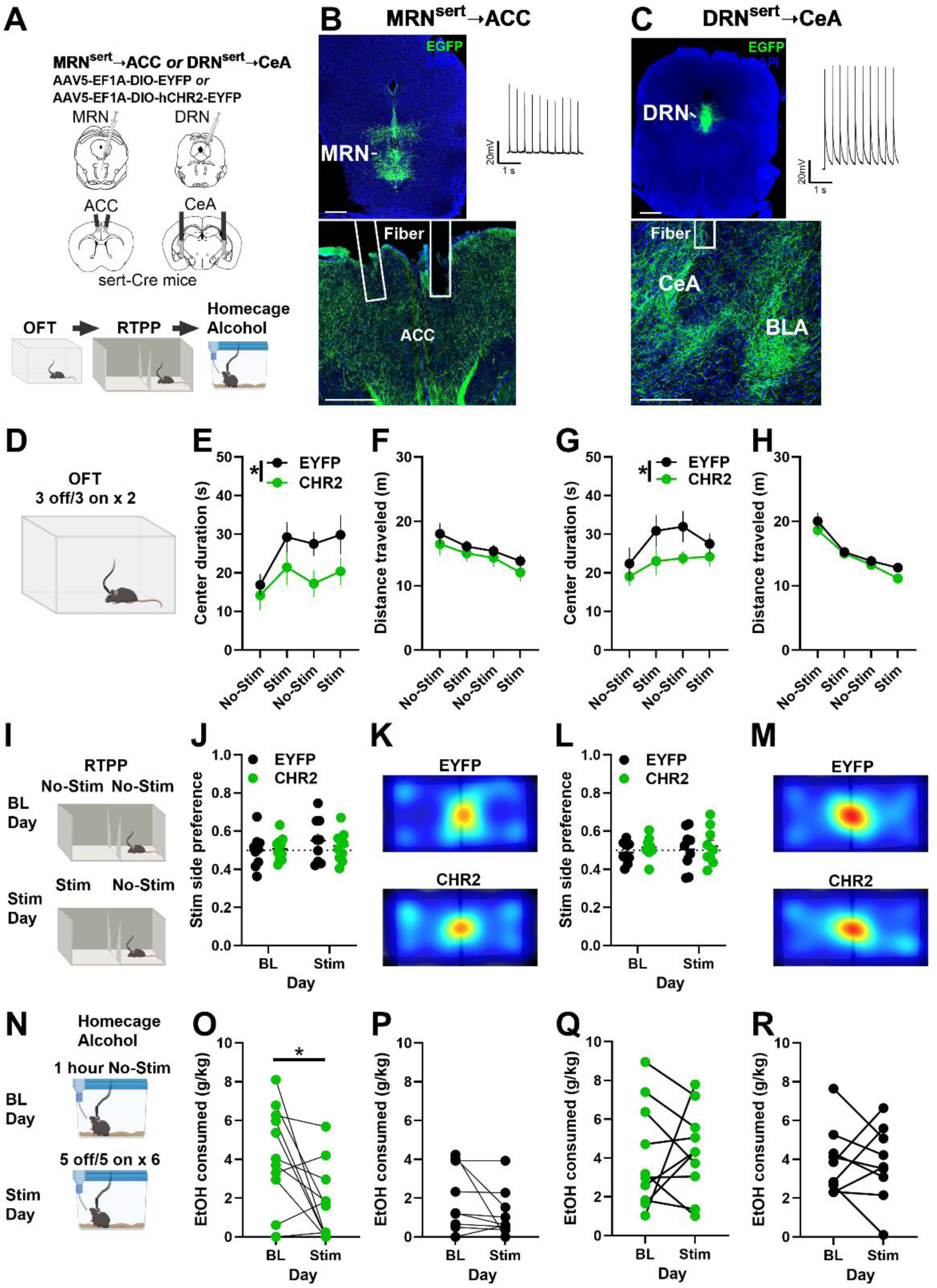
5HT terminal stimulation in the ACC and CeA increases avoidance behavior, but only stimulation in the ACC decreased alcohol intake. **(A)** Experimental timeline. Male and female Sert-cre mice received AAV5-EF1A-DIO-EYFP or AAV5-EF1A-DIO-hCHR2-EYFP in the MRN or DRN and bilateral fiber optic implants in the anterior cingulate cortex (ACC) or central nucleus of the amygdala (CeA) to allow for circuit specific manipulation of MRN^Sert^→ACC or DRN^Sert^→CeA, respectively. They were subsequently tested in the open field test (OFT), real-time place preference (RTPP), and home cage alcohol drinking assays. All data analyzed collapsed across sex due to lack of clear sex effects. **(B, upper left)** Representative image of EYFP expression in the MRN and **(B, upper right)** optically evoked current from a MRN^Sert^ neuron. Scale bars represent 1 mm. **(B, bottom)** EYFP expression in MRN^Sert^ processes and fiber implants in ACC. **(C, upper left)** Representative image of EYFP expression in the DRN and **(C, upper right)** optically evoked current from a DRN^Sert^ neuron. **(C, bottom)** Representative image of EYFP expression in DRN^Sert^ processes and fiber implant in the CeA. **(D)** Mice were allowed to explore an open field for 12 minutes. Laser stimulation was applied at 20 hz for 3-minute blocks during the session (3 min off, 3 min on, x2). **(E)** Center duration shown by 3-minute bin across session for MRN^Sert^→ACC mice. CHR2 mice had lower center duration than EYFP mice (2-way ANOVA, group main effect, F_(1,72)_ = 7.59, *p* = 0.007) and center duration changed across time (time main effect, F_(3,72)_ = 2.75, p = 0.05). **(F)** Distance traveled during the OFT. **(G)** Center duration shown by 3-minute bin across session for DRN^Sert^→CeA mice. CHR2 mice had lower center duration than EYFP mice (2-way ANOVA, group main effect, F_(1,64)_ = 6.07, *p* = 0.02). **(H)** Distance traveled during the OFT. **(I)** RTPP experiment design. Mice were allowed to freely explore a two-chamber apparatus for 20 minutes with no stimulation (No-Stim) on either side (baseline, BL). The following day, mice were allowed to explore the apparatus but entry into one of the sides triggered 20 hz stimulation (Stim). **(J)** Quantification of preference for the stimulation side on BL and Stim day for MRN^Sert^→ACC mice. **(K)** Representative heatmaps of time spent in the apparatus from EYFP and CHR2 mice. **(L)** Quantification of preference for the stimulation side on BL and Stim day for DRN^Sert^→CeA mice. **(M)** Representative heatmaps of time spent in the apparatus from EYFP and CHR2 mice. **(N)** Home cage alcohol drinking experiment design. Mice were acclimated to drinking 20% alcohol for four 1-hour sessions prior to BL day. On BL day, mice were allowed to drink 20% alcohol for 1 hour while tethered without laser stimulation. On Stim day, mice were allowed to drink 20% alcohol for 1 hour and received 20hz laser stimulation in 5-minute blocks (5 off/5 on, x6). BL and Stim days were in counterbalanced order. **(O)** Alcohol consumed during the BL and Stim sessions of CHR2 MRN^Sert^→ACC mice. Mice consumed less alcohol on stim day than at BL (Welch’s t-test, t_(18.74)_ = 2.72, p = 0.01). **(P)** Alcohol consumed during the BL and Stim sessions of EYFP MRN^Sert^→ACC mice. **(Q)** Alcohol consumed during the BL and Stim sessions of CHR2 DRN^Sert^→CeA mice. **(R)** Alcohol consumed during the BL and Stim sessions of EYFP DRN^Sert^→CeA mice.

## Discussion

5HT signaling is critical for healthy brain function and becomes dysregulated in AUD and comorbid psychopathologies^6^, yet there is a limited understanding of region-specific 5HT dynamics. The ACC and CeA play distinct roles in the progression of AUD and have opposite changes in SERT expression in problem drinkers^7,9^, suggesting that alcohol misuse may elicit region-specific changes in 5HT signaling. However, the extent to which *in vivo* 5HT dynamics in the ACC and CeA are altered by binge alcohol consumption is poorly understood. We aimed to address this gap in knowledge using *in vivo* fiber photometry recordings of the fluorescent 5HT biosensor, GRAB5HT, in alcohol-naïve mice and after 1 and 4 weeks of DID. This time course was chosen based on prior evidence of dysfunction in ACC neuronal excitability following similar periods of DID^3^. We found DID elicited sex-, brain region-, and exposure-dependent alterations in 5HT dynamics. The consumption of alcohol, sucrose, and HFD elicited opposing responses in 5HT in the ACC and CeA, which exhibited decreased and increased signal, respectively, and early binge drinking blunted ACC and enhanced CeA 5HT dynamics after 1 week of drinking experience. In contrast, aversive and anxiogenic stimuli increased 5HT in both regions, and 4 weeks of DID produced sex-specific changes in these responses that were modality dependent. Finally, genetically targeted optogenetic stimulation of 5HT terminals in the ACC and CeA similarly elicited an anxiety-like state, while only stimulation in the ACC blunted alcohol consumption. Together, these data provide insight into the sex- and region-specific impact of binge drinking on 5HT dynamics.

Consummatory behaviors evoked opposing 5HT signaling in the ACC and CeA. This is in line with prior investigations into the primary 5HT inputs of these structures, the MRN and DRN, respectively. Specifically, MRN 5HT neurons are suppressed^29^, while DRN neurons are activated^30^, during reward consumption. The opposing reward-associated 5HT signals likely reflect the distinct roles the downstream structures play in reward processing and action selection. The ACC provides top-down gating of goal-directed behaviors based on performance demands without encoding an explicit reward value^31,32^. *In vitro*, application of 5HT increases action potential firing of cortical fast spiking interneurons and decreases firing in pyramidal neurons^33^. Though pyramidal neurons express Gq-coupled 5HT2a, which can increase spontaneous excitatory post-synaptic currents *in vitro* through non-synchronous glutamate release^34^, *in vivo* agonism of 5HT2a in frontal cortex blunts pyramidal firing rate^35^. Together with our data, these observations suggest that 5HT suppression of ACC output is reduced during consummatory behaviors, which may serve to promote task attention and increase intake^36,37^.

ACC 5HT signaling was blunted during alcohol consumption after one week of DID similarly in males and females. This effect contrasts with the sex-specific effects of DID on 5HT dynamics in response to other consummatory behaviors. Curiously, the 5HT response to alcohol returned after 4 weeks of DID. Thus, DID elicits dysfunction in the ACC specific to alcohol that varies with the length of alcohol history, in line with prior literature showing altered ACC pyramidal neuron excitability after 1 week of binge drinking that normalized by 4 weeks^3^. In our earlier work, alcohol consumption elicited a reduction of 5HT on to 5HT2C-R containing neurons in the bed nucleus of the stria terminalis (BNST)^26^. This is similar to what we observed in the ACC, suggesting the BNST 5HT signal may be MRN biased. The ACC is critical for decision making and risk assessment, as abstinent individuals with a history of heavy alcohol use exhibit higher risky decision making and diminished ACC recruitment relative to healthy controls^10^. Importantly, propensity for risk taking is a strong predictor of continued alcohol use^38^. Thus, the blunting of 5HT dynamics during early episodes of binge drinking could underly ACC dysfunction that shapes future risky decision making, potentially heightening the risk for the development of an AUD.

In contrast, CeA neurons signal both valence and salience, with approximately equal subpopulations encoding positive and negative valence^39^. Prior to alcohol exposure, 5HT abolishes CeA neuronal firing by enhancing presynaptic GABA release in a 5HT2C-dependent manner^40^. In alcohol dependence and during withdrawal, CeA 5HT sensitivity is blunted^15^. Our observation that DID males show enhanced 5HT response to alcohol consumption suggests that binge drinking enhances the alcohol reward or salience signal in the CeA. Alternatively, the increased 5HT signal may be a compensatory response to account for blunted 5HT sensitivity in the CeA. Prior work did not include female subjects, so it is unclear how alcohol alters CeA 5HT function in females, making this an important topic for future investigation.

We were surprised to find that male water mice showed a nearly complete loss of 5HT response in the CeA to sucrose consumption across recording sessions that was preserved in DID mice, with females showing no such adaptation in signal. In men, self-reported liking of high concentration sucrose is higher in those with a history of AUD without altering the ability to discriminate between sucrose concentrations^41^, supporting the idea that sucrose preference may be predictive of AUD risk^42^. Notably, women with AUD do not exhibit increased sweet liking^43^. One potential interpretation of our data is that binge drinking may elicit increased salience or reward value for sweet substances which in turn maintains the reward-like 5HT response in the CeA. These data are interesting in light of contrasting clinical evidence that alcohol use severity is predictive of increased incentive motivation value of alcohol cues relative to natural reward cues^44^. Thus, alcohol may elicit sex-dependent dissociable effects on conditioned and unconditioned stimuli (i.e., alcohol cues versus alcohol itself).

In contrast to the 5HT dynamics during reward, 5HT in the ACC and CeA increased during aversive and anxiogenic conditions, consistent with threat and aversion encoding by MRN and DRN 5HT neurons^29,45^. Interestingly, DID history did not uniformly disrupt the 5HT response to aversive stimuli. Instead, it produced discrete sex-, brain region-, and stimulus modality-specific effects. In males, DID produced elevated ACC 5HT signal during explore/retreat action selection in the EPM. Given that the probability to explore decreased with higher ACC 5HT signal, increased ACC 5HT may have contributed to the decrease in open arm entries exhibited by males. Females by comparison showed altered 5HT during sensory processing and center exploration without altering anxiety-like behavior in the EPM. One potential limitation of the present study is that the effects of DID on anxiety-like behavior were assessed at only one time point. A more protracted withdrawal period may have revealed behavioral effects in females, as 2-4 weeks of abstinence from DID results in female-specific behavioral disinhibition^46^. Another potential limitation is that our approach is not optimized for changes in baseline 5HT levels. Together, the observed pattern of effects on aversive processing suggests that DID does not uniformly disrupt 5HT signaling and highlights the need for further investigation into the circuit-selective impact of alcohol misuse.

To understand how manipulation of 5HT signaling in the CeA and ACC contributes to changes in behavior, we used an optogenetic approach. We first performed anatomical experiments to confirm the most abundant 5HT innervation of each region, with the ACC receiving predominantly MRN input, and the CeA receiving predominantly DRN input. Surprisingly, 5HT terminal stimulation in both the ACC and CeA was anxiogenic in the open field test but failed to alter place preference. This suggests that 5HT signal in these structures is not inherently positive or negative but may serve to modulate the behavioral response to valanced sensory input, perhaps as a metric of salience, in line with the established neuromodulatory role of 5HT. In addition, we found that stimulation of 5HT terminals in the ACC led to a reduction in alcohol drinking. Given that 5HT signal decreased in the ACC with both alcohol and sucrose drinking, and we did not examine the impact of optostimulation on sucrose consumption, it is unclear if the effect of terminal stimulation is a general suppression of consummatory behavior or an alcohol specific effect. However, this finding aligns with the idea that high levels of 5HT release in the ACC can serve to inhibit alcohol consumption.

Alcohol misuse has a lasting impact on the 5HT system, yet our exploration of brain region- and sex-specific effects of binge alcohol use on real-time 5HT dynamics has only recently begun^26^. Our data indicate that alcohol elicits discrete region-specific remodeling of 5HT signaling with functional implications for reward, stress and anxiety, and action selection. Given that 5HT dysfunction was especially evident after 1 week of DID, early experiences with alcohol misuse may represent a window of vulnerability in the progression of AUD. Together, these data build on a very limited literature characterizing the effects of binge drinking on 5HT dynamics and suggest a potential mechanism for targeted intervention.

## Works Cited

1. Paljärvi, T. et al. Binge drinking and depressive symptoms: a 5-year population-based cohort study. Addiction 104, 1168–1178 (2009).

2. Jennison, K. M. The Short-Term Effects and Unintended Long-Term Consequences of Binge Drinking in College: A 10-Year Follow-Up Study. Am. J. Drug Alcohol Abuse 30, 659–684 (2004).

3. Cannady, R., Nimitvilai-Roberts, S., Jennings, S. D., Woodward, J. J. & Mulholland, P. J. Distinct Region- and Time-Dependent Functional Cortical Adaptations in C57BL/6J Mice after Short and Prolonged Alcohol Drinking. eneuro 7, ENEURO.0077-20.2020 (2020).

4. Belmer, A., Depoortere, R., Beecher, K., Newman-Tancredi, A. & Bartlett, S. E. Neural serotonergic circuits for controlling long-term voluntary alcohol consumption in mice. Mol. Psychiatry 27, 4599–4610 (2022).

5. Lee, K. M., Coehlo, M., McGregor, H. A., Waltermire, R. S. & Szumlinski, K. K. Binge alcohol drinking elicits persistent negative affect in mice. Behav. Brain Res. 291, 385–398 (2015).

6. Bach, H. et al. Alcoholics Have More Tryptophan Hydroxylase 2 mRNA and Protein in the Dorsal and Median Raphe Nuclei. Alcohol. Clin. Exp. Res. 38, 1894–1901 (2014).

7. Mantere, T. et al. Serotonin Transporter Distribution and Density in the Cerebral Cortex of Alcoholic and Nonalcoholic Comparison Subjects: A Whole-Hemisphere Autoradiography Study. Am. J. Psychiatry 159, 599–606 (2002).

8. Storvik, M., Hakkinen, M., Tupala, E. & Tiihonen, J. 5-HT1A Receptors in the Frontal Cortical Brain Areas in Cloninger Type 1 and 2 Alcoholics Measured by Whole-Hemisphere Autoradiography. Alcohol Alcohol 44, 2–7 (2008).

9. Storvik, M., Tiihonen, J., Haukijärvi, T. & Tupala, E. Amygdala serotonin transporters in alcoholics measured by whole hemisphere autoradiography. Synapse 61, 629–636 (2007).

10. Claus, E. D. & Hutchison, K. E. Neural Mechanisms of Risk Taking and Relationships with Hazardous Drinking. Alcohol. Clin. Exp. Res. 36, 932–940 (2012).

11. Zakiniaeiz, Y., Scheinost, D., Seo, D., Sinha, R. & Constable, R. T. Cingulate cortex functional connectivity predicts future relapse in alcohol dependent individuals. NeuroImage Clin. 13, 181–187 (2017).

12. Cannady, R., Nimitvilai-Roberts, S., Jennings, S. D., Woodward, J. J. & Mulholland, P. J. Distinct Region- and Time-Dependent Functional Cortical Adaptations in C57BL/6J Mice after Short and Prolonged Alcohol Drinking. eneuro 7, ENEURO.0077-20.2020 (2020).

13. Roberto, M., Kirson, D. & Khom, S. The Role of the Central Amygdala in Alcohol Dependence. Cold Spring Harb. Perspect. Med. 11, a039339 (2021).

14. Yoshimoto, K. et al. Alcohol enhances characteristic releases of dopamine and serotonin in the central nucleus of the amygdala. Neurochem. Int. 37, 369–376 (2000).

15. Khom, S. et al. Alcohol Dependence and Withdrawal Impair Serotonergic Regulation of GABA Transmission in the Rat Central Nucleus of the Amygdala. J. Neurosci. 40, 6842–6853 (2020).

16. Marcinkiewcz, C. A. et al. Serotonin engages an anxiety and fear-promoting circuit in the extended amygdala. Nature 537, 97–101 (2016).

17. Roland, A. V. et al. Acute and chronic alcohol modulation of extended amygdala calcium dynamics. Alcohol 116, 53–64 (2024).

18. Flanigan, M. E. et al. Subcortical serotonin 5HT(2c) receptor-containing neurons sex-specifically regulate binge-like alcohol consumption, social, and arousal behaviors in mice. Nat Commun 14, 1800 (2023).

19. Mathis, A. et al. DeepLabCut: markerless pose estimation of user-defined body parts with deep learning. Nat. Neurosci. 21, 1281–1289 (2018).

20. Goodwin, N. L. et al. Simple Behavioral Analysis (SimBA) as a platform for explainable machine learning in behavioral neuroscience. Nat. Neurosci. 27, 1411–1424 (2024).

21. Neira, S. et al. Chronic alcohol consumption alters home-cage behaviors and responses to ethologically relevant predator tasks in mice. Alcohol. Clin. Exp. Res. 46, 1616–1629 (2022).

22. Schindelin, J., et al. Fiji: an open-source platform for biological-image analysis. Nat. Methods 9, 676–682 (2012).

23. Marcinkiewcz, C. A. et al. Serotonin engages an anxiety and fear-promoting circuit in the extended amygdala. Nature 537, 97–101 (2016).

24. Olivier, J. D. A. et al. A study in male and female 5-HT transporter knockout rats: An animal model for anxiety and depression disorders. Neuroscience 152, 573–584 (2008).

25. Jacobs, B. L. & Cohen, A. Differential Behavioral Effects of Lesions of the Median or Dorsal Raphe Nuclei in Rats: Open Field and Pain-Elicited Aggression.

26. Flanigan, M. E. et al. Subcortical serotonin 5HT2c receptor-containing neurons sex-specifically regulate binge-like alcohol consumption, social, and arousal behaviors in mice. Nat. Commun. 14, 1800 (2023).

27. Ren, J. et al. Single-cell transcriptomes and whole-brain projections of serotonin neurons in the mouse dorsal and median raphe nuclei. eLife 8, e49424 (2019).

28. Muzerelle, A., Scotto-Lomassese, S., Bernard, J. F., Soiza-Reilly, M. & Gaspar, P. Conditional anterograde tracing reveals distinct targeting of individual serotonin cell groups (B5–B9) to the forebrain and brainstem. Brain Struct. Funct. 221, 535–561 (2016).

29. Kawai, H. et al. Median raphe serotonergic neurons projecting to the interpeduncular nucleus control preference and aversion. Nat. Commun. 13, 7708 (2022).

30. Li, Y. et al. Serotonin neurons in the dorsal raphe nucleus encode reward signals. Nat. Commun. 7, 10503 (2016).

31. Kim, J.-H., Ma, D.-H., Jung, E., Choi, I. & Lee, S.-H. Gated feedforward inhibition in the frontal cortex releases goal-directed action. Nat. Neurosci. 24, 1452–1464 (2021).

32. Huda, R. et al. Distinct prefrontal top-down circuits differentially modulate sensorimotor behavior. Nat. Commun. 11, 6007 (2020).

33. Zhong, P. & Yan, Z. Differential Regulation of the Excitability of Prefrontal Cortical Fast-Spiking Interneurons and Pyramidal Neurons by Serotonin and Fluoxetine. PLoS ONE 6, e16970 (2011).

34. Aghajanian, G. K. & Marek, G. J. Serotonin, via 5-HT receptors, increases EPSCs in layer V pyramidal cells 2A of prefrontal cortex by an asynchronous mode of glutamate release.

35. Wang, S. et al. In vivo effects of activation and blockade of 5-HT2A/2C receptors in the firing activity of pyramidal neurons of medial prefrontal cortex in a rodent model of Parkinson’s disease. Exp. Neurol. 219, 239–248 (2009).

36. Koike, H. et al. Chemogenetic Inactivation of Dorsal Anterior Cingulate Cortex Neurons Disrupts Attentional Behavior in Mouse. Neuropsychopharmacology 41, 1014–1023 (2016).

37. Yu, Y. H. et al. Optogenetic stimulation in the medial prefrontal cortex modulates stimulus valence from rewarding and aversive to neutral states. Front. Psychiatry 14, 1119803 (2023).

38. MacPherson, L., Magidson, J. F., Reynolds, E. K., Kahler, C. W. & Lejuez, C. W. Changes in Sensation Seeking and Risk-Taking Propensity Predict Increases in Alcohol Use Among Early Adolescents. Alcohol. Clin. Exp. Res. 34, 1400–1408 (2010).

39. Kong, M.-S., Ancell, E., Witten, D. M. & Zweifel, L. S. Valence and salience encoding in the central amygdala. eLife 13, RP101980 (2025).

40. Hon, O. J. et al. Serotonin modulates an inhibitory input to the central amygdala from the ventral periaqueductal gray. Neuropsychopharmacology 47, 2194–2204 (2022).

41. Evidence of preference for a high-concentration sucrose solution in alcoholic men. Am. J. Psychiatry 154, 269–270 (1997).

42. Bouhlal, S., Farokhnia, M., Lee, M. R., Akhlaghi, F. & Leggio, L. Identifying and Characterizing Subpopulations of Heavy Alcohol Drinkers Via a Sucrose Preference Test: A Sweet Road to a Better Phenotypic Characterization? Alcohol Alcohol 53, 560–569 (2018).

43. Lange, L. A., Kampov-Polevoy, A. B. & Garbutt, J. C. Sweet Liking and High Novelty Seeking: Independent Phenotypes Associated with Alcohol-related Problems. Alcohol Alcohol 45, 431–436 (2010).

44. Martins, J. S. et al. Differential brain responses to alcohol-related and natural rewards are associated with alcohol use and problems: Evidence for reward dysregulation. Addict. Biol. 27, e13118 (2022).

45. Paquelet, G. E. et al. Single-cell activity and network properties of dorsal raphe nucleus serotonin neurons during emotionally salient behaviors. Neuron 110, 2664–2679.e8 (2022).

46. Rivera-Irizarry, J. K. et al. Sex differences in binge alcohol drinking and the behavioral consequences of protracted abstinence in C57BL/6J mice. Biol. Sex Differ. 14, 83 (2023).

