## Supplemental Data for "Selective dysregulation of serotonin dynamics in the anterior cingulate cortex and central amygdala following binge alcohol consumption"

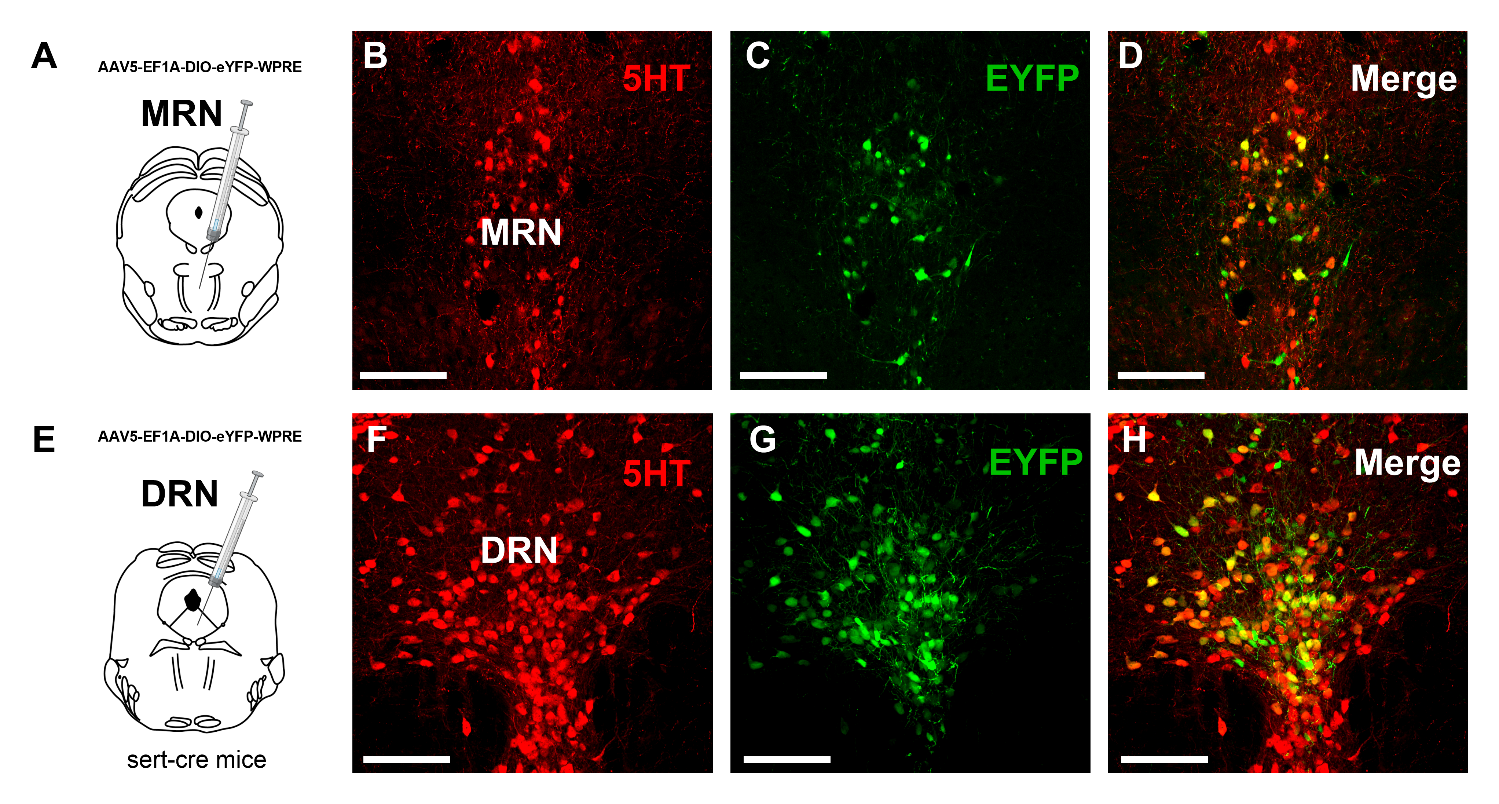

**Supplemental Figure 1. Immunohistochemical validation of Sert-cre targeting of 5HT expressing neurons.** Sert-cre mice received a cre-dependent EYFP virus in the median or dorsal raphe nuclei (MRN, DRN, respectively). **(A)** Viral targeting and immunohistochemical labeling of **(B)** 5HT (red) and **(C)** EYFP in the MRN. **(D)** Merged image of the MRN. **(E)** Viral targeting and immunohistochemical labeling of **(F)** 5HT (red) and **(G)** EYFP in the DRN. **(H)** Merged image of the DRN. Scale bar equivalent to 1 mm.

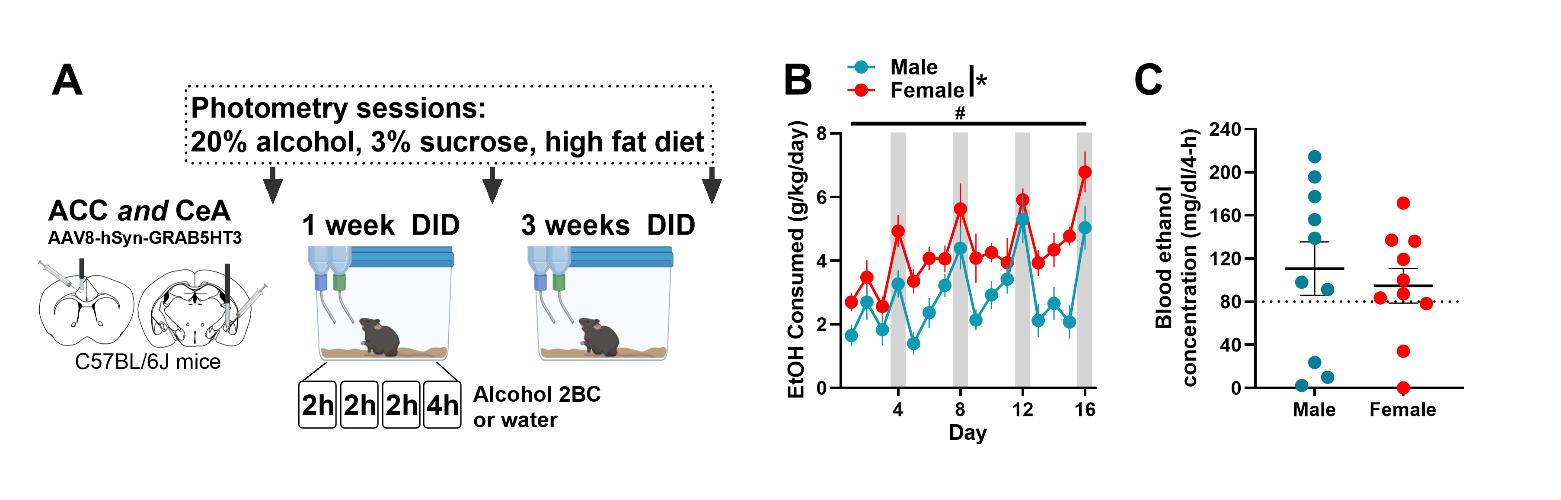

**Supplemental Figure 2. Drinking history and blood alcohol concentration during 4 weeks of Drinking in the Dark binge drinking. (A)** Experimental design. Male and female C57BL/6J mice received GRAB5HT and fiber optic implants directed at anterior cingulate cortex and central nucleus of the amygdala and underwent 4 weeks of 2-bottle choice Drinking in the Dark (DID) binge drinking of 20% alcohol (N = 10 male, 10 female) or served as water controls (N = 10 male, 9 female). Photometry recording sessions to assess *in vivo* 5HT dynamics while consuming 20% alcohol, 3% sucrose, and high fat diet (HFD) were conducted in alcohol naïve mice and after 1 and 4 weeks of DID. **(B)** Alcohol consumption across 4-weeks of DID sessions normalized to body weight increased across time and females drank more than males (2-way ANOVA, day main effect, F_(15,288)_ = 9.30, *p* < 0.0001, Sidak’s post hoc, day 4 < day 16, p = 0.04; Sex main effect, F_(1,288)_ = 60.04, *p* < 0.001). **(C)** Blood alcohol concentration taken immediately following the final DID session. The dotted line at 80 mg/dl represents binge level of intoxication. Data are represented as mean±SEM.

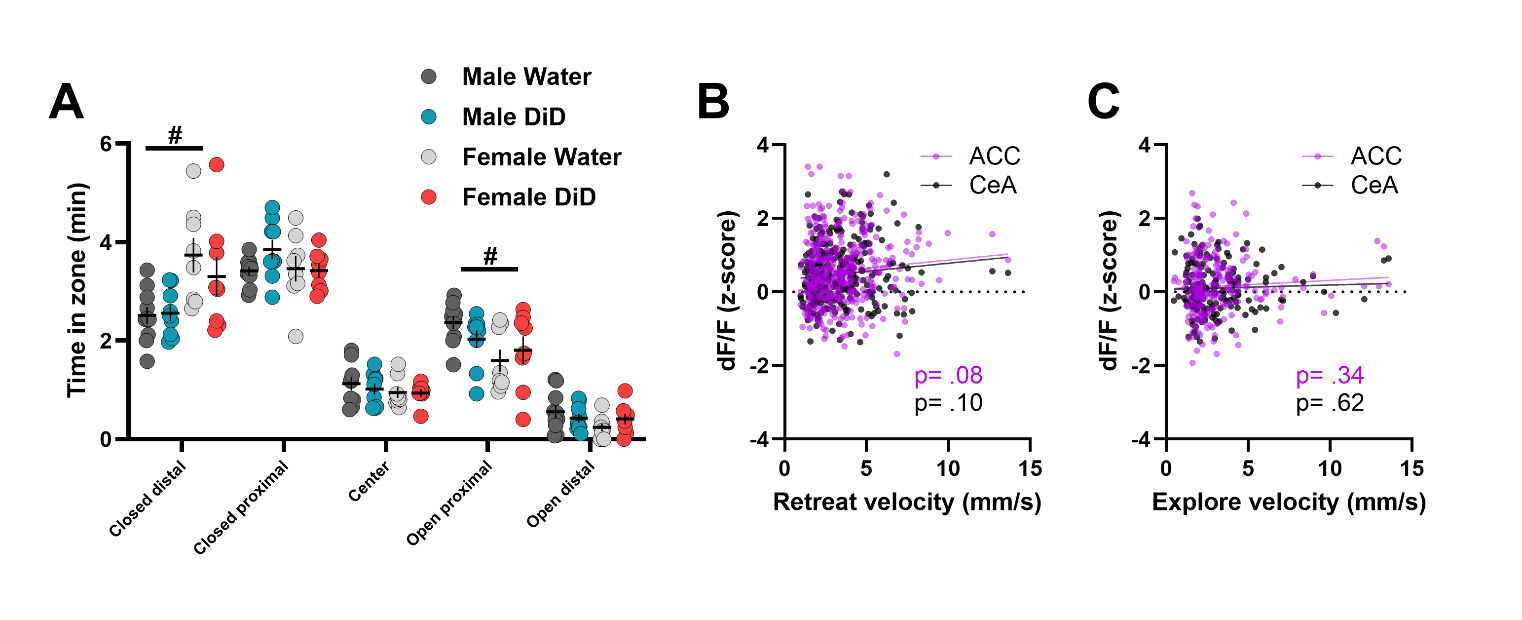

**Supplemental Figure 3. Time spent zones of elevated plus maze and relationship between 5HT signal and velocity of exploration and retreat. (A)** Time spent in each zone of the elevated plus maze. Females spent more time in the distal portion of the closed arm and less time in the proximal part of the open arm than males (3-way ANOVA, sex x zone interaction, F_(4,124)_ = 7.64, *p* < 0.001, closed distal male vs female *p* = 0.001, open proximal male vs female *p* = 0.02). **(B)** Scatterplot of retreat velocity relative to associated 5HT signal in the ACC and CeA. No significant correlation was observed (Pearson’s correlation, ACC r = 0.09, *p* = 0.08, CeA r = 0.10, *p* = 0.10). **(C)** Scatterplot of explore velocity and 5HT signal in the ACC and CeA. No significant correlation was observed (Pearson’s correlation, ACC r = 0.06, *p* = 0.34, CeA r = 0.04, *p* = 0.62). Data are represented as mean±SEM.

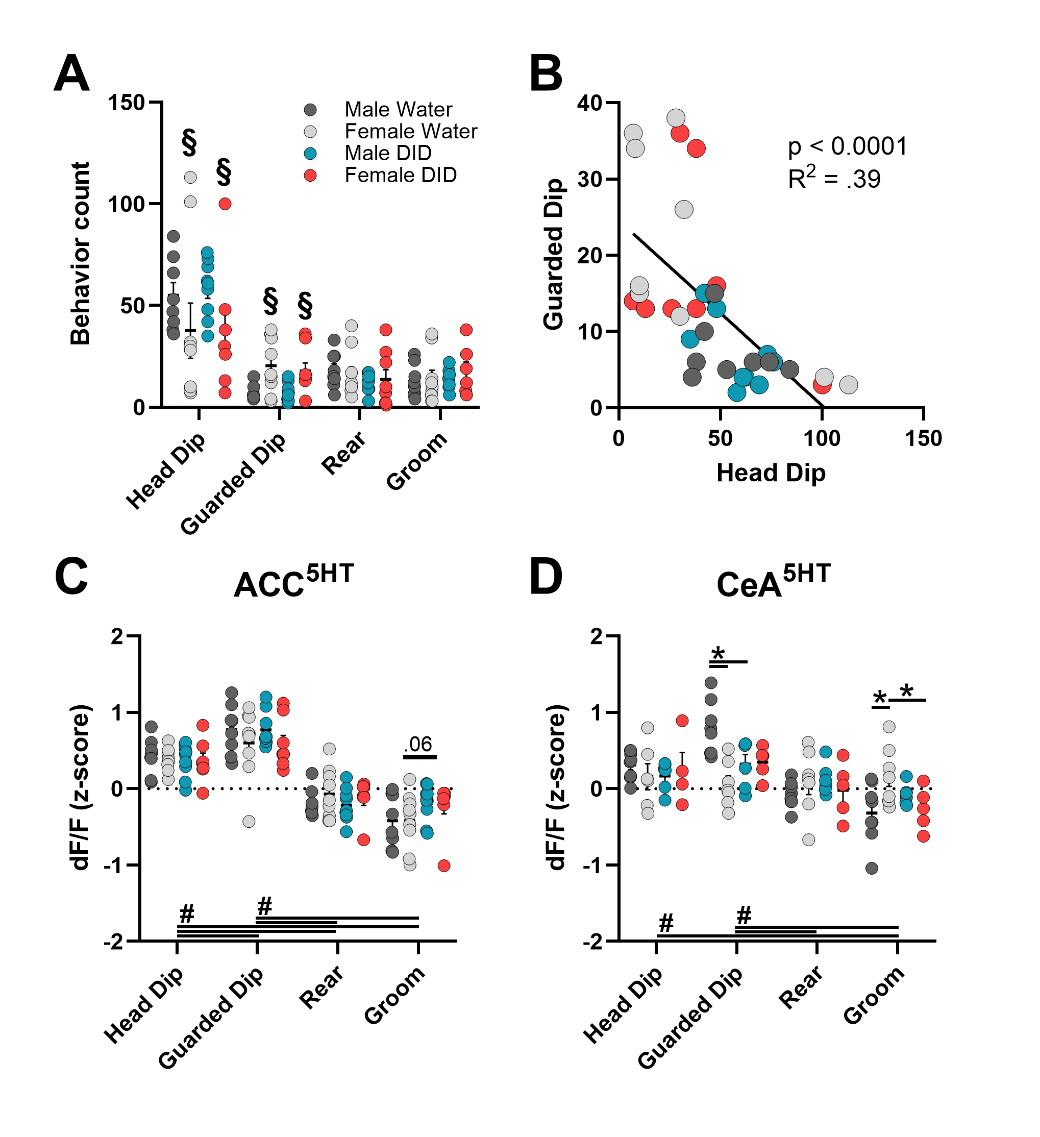

**Supplemental Figure 4. Sex-dependent effects of DID on 5HT signaling during threat assessment and stress coping.** Male and female C57BL/6J mice received GRAB5HT and fiber optic implants directed at anterior cingulate cortex and central nucleus of the amygdala and underwent 4 weeks of 2-bottle choice Drinking in the Dark (DID) binge drinking of 20% alcohol (N = 10 male, 10 female) or served as water controls (N = 10 male, 9 female). Mice were then allowed to explore an elevated plus maze (EPM) for 10 minutes. Behavior was recorded from above and scored using DeepLabCut and SiMBA machine learning software. **(A)** Females exhibited less head dips and more guarded head dips than males, with no effect of DID on behavior counts (head dips, 2 x 2 ANOVA, sex main effect only, F_(1,30)_ = 4.05, *p* = 0.05; guarded dip, 2 x 2 ANOVA, sex main effect only, F_(1,30)_ = 14.07, *p* < 0.001; All DID effects *p* > 0.05). **(B)** The number of head dips negatively correlated with guarded dips (Pearson’s correlation, r = -0.63, *p* < 0.001). **(C)** Z-scored GRAB5HT signal in the ACC during head dip, guarded dip, rear, and groom in the EPM averaged across bouts. There were no significant sex or group effects on 5HT signal during EPM behaviors, but there was a trend towards a blunted decrease in groom signal in DID mice (2 x 2 ANOVA, F_(1,30)_ = 3.88, *p* = 0.06). **(D)** Z-scored GRAB5HT signal in the CeA during head dip, guarded dip, rear, and groom in the EPM averaged across bouts. DID blunted 5HT response during guarded dip in male mice and female water mice had a lower 5HT response than male water mice (2 x 2 ANOVA, sex x group interaction, F_(1,22)_ = 9.78, *p* = 0.005, Sidak’s post hoc tests, male water > male DID, *p* = 0.009, male water > female water, *p* < 0.001). In contrast, DID blunted 5HT response during grooming in females and female water mice had a higher 5HT response than male water mice (2 x 2 ANOVA, sex x group interaction, F_(1,22)_ = 6.78, *p* = 0.02, Sidak’s post hoc tests, female water > female DID, *p* = 0.03, male water < female water, *p* = 0.008). Data are represented as mean±SEM.

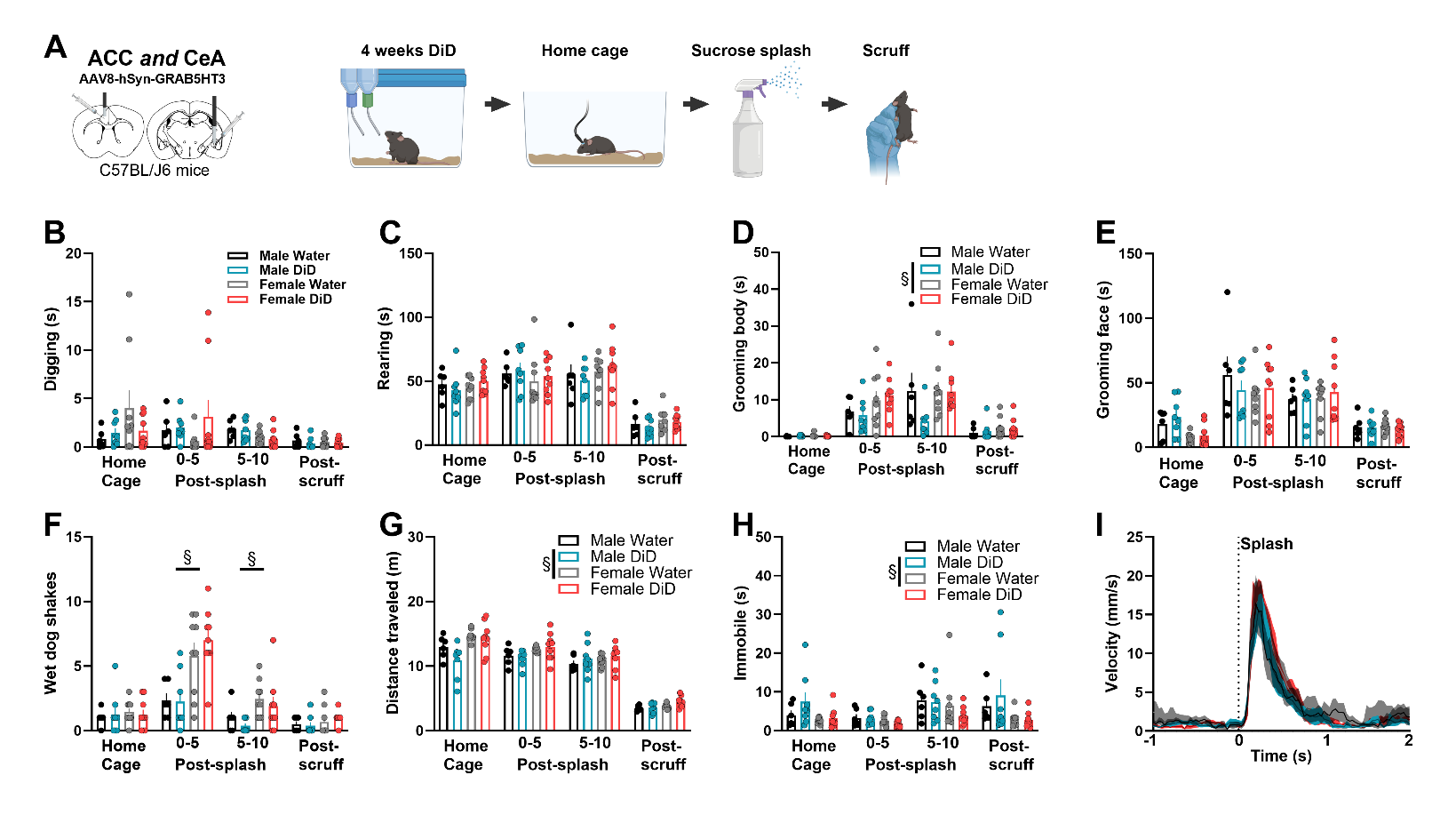

**Supplemental Figure 5. No effect of DID on home cage behaviors during the sucrose splash and scruff assay.** **(A)** Experimental timeline. Male and female C57BL/6J mice received GRAB5HT and fiber optic implants directed at anterior cingulate cortex and central nucleus of the amygdala and underwent 4 weeks of 2-bottle choice Drinking in the Dark (DID) binge drinking of 20% alcohol (N = 10 male, 10 female) or served as water controls (N = 10 male, 9 female). Mice were allowed to explore their home cage for 5 minutes then received a spray of 10% sucrose solution directed at their back. Ten minutes later mice were scruffed for 1 minute and then allowed to explore their home cage for 2 minutes. Behaviors were analyzed from an overhead camera using DLC and SiMBA. The behaviors analyzed included **(B)** digging, **(C)** rearing, **(D)** grooming body, **(E)** grooming face, **(F)** wet dog shake, **(G)** distance traveled, **(H)** immobility, and **(I)** post-splash velocity. Females exhibited more wet dog shakes post-splash (2 x 2 x 2 ANOVA, sex x time interaction, F(2,56) = 25.11, *p* < 0.001, Male < Female, *p* < 0.05), more overall grooming body (2 x 2 x 2 ANOVA, sex effect, F(1,28) = 9.56, *p* = 0.004), more distance traveled (2 x 2 x 2 ANOVA, sex effect, F(1,28) = 7.77, *p* = 0.009; group x time interaction, F(3,84) = 5.62, *p* = 0.001, Sidak’s post hoc tests *p* > 0.05), and spent less time immobile (2 x 2 x 2 ANOVA, sex effect, F(1,28) = 6.60, *p* = 0.02). Data are represented as mean±SEM. ^&^significant effect of sex.
